# Effects of *Wolbachia* on transposable element activity largely depend on *Drosophila melanogaster* host genotype

**DOI:** 10.1101/2022.07.21.500779

**Authors:** Ana T. Eugénio, Marta S. P. Marialva, Patrícia Beldade

## Abstract

Transposable elements (TEs) are repetitive DNA sequences capable of changing position in host genomes, thereby causing mutations. TE insertions typically have deleterious effects but they can also be beneficial. Increasing evidence of the contribution of TEs to adaptive evolution further raises interest in understanding what factors impact TE activity. Based on previous studies associating the bacterial endosymbiont *Wolbachia* to changes in the abundance of piRNAs, a mechanism for TE repression, and to transposition of specific TEs, we hypothesized that *Wolbachia* infection would interfere with TE activity. We tested this hypothesis by studying expression of 14 TEs in a panel of 25 *Drosophila melanogaster* host genotypes, naturally infected with *Wolbachia* and annotated for TE insertions. The host genotypes differed significantly in *Wolbachia* titers inside individual flies, with broad-sense heritability around 20%, and in the number of TE insertions, which depended greatly on TE identity. By removing *Wolbachia* from the target host genotypes, we generated a panel of 25 pairs of *Wolbachia*-positive and *Wolbachia*-negative lines in which we quantified transcription levels our target TEs. We found variation in TE expression that was dependent on *Wolbachia* status, TE identity, and host genotype. Comparing between pairs of *Wolbachia*-positive and *Wolbachia*-negative flies, we found that *Wolbachia* removal affected TE expression in 23.7% of the TE-genotype combinations tested, with up to 4.6 times differences in median level of transcript. Our data shows that *Wolbachia* can impact TE activity in host genomes, underscoring the importance this endosymbiont can have in the generation of genetic novelty in hosts.

## Introduction

Transposable elements (TEs) are repetitive DNA sequences capable of changing position independently in the host genome (Bourque et al. 2018; Mérel et al. 2020), and make up for a significant fraction of many eukaryotic genomes (Guio and González 2019). They are divided into two major classes, depending on whether the mechanism of transposition does (for retrotransposons) or does not (for DNA transposons) involve an RNA intermediate that is reverse-transcribed before integrating back into the host genome (Bourque et al. 2018). TE insertions can cause mutations, which typically have deleterious effects because they disrupt proper gene function in a variety of manners (McFaddenaf et al. 1997; Hedges et al. 2007; Belancio et al. 2008; Ayarpadikannan and Kim 2014). Consequently, host organisms have evolved mechanisms to control and repress TE activity, including the germline-specific piRNA pathway in animals (Tóth et al. 2016). On the other hand, an increasing number of studies have been providing compelling examples of TE insertions with positive effects on host fitness, contributing to adaptation (González et al. 2009; González et al. 2010; van’t Hof et al. 2016), stress resistance (Guio et al. 2014; Pereira and Ryan 2019), and the origin of novel traits (Emera et al. 2012; Bennetzen and Wang 2014; Santos et al. 2014, Trizzino et al. 2017). Moreover, TEs might also contribute to reproductive isolation, as in the case of TE-mediated hybrid incompatibility (Petrov et al. 1995; Serrato-Capuchina et al. 2018). TE contribution to adaptive evolution and diversification raises interest in understanding what factors impact TE activity.

TE activity differs between TEs (Venner et al. 2009; Mérel et al. 2020) and between host genotypes (Anderson et al. 2019; Signor 2020). Furthermore, studies on different organisms have shown that TE activity can be affected by external environmental factors, including temperature (Chen et al. 2018), radiation (Newman et al. 2014), heavy metals (Habibi et al. 2014), starvation (Rep et al. 2005), and various other stressors (Miousse et al. 2015). Not much is known about how these factors can affect the molecular mechanisms responsible for TE regulation, including the piRNA pathway. On the other hand, *Wolbachia*, a common endosymbiotic bacterium, has been shown to affect piRNA abundance (Mayoral et al. 2014) and the rate of transposition of the retrotransposon gypsy (Touret et al. 2014). Moreover, the invasion of the DNA transposon P-element in populations of *Drosophila* reportedly co-occurred with a replacement of *Wolbachia* strain infecting those flies (Riegler et al. 2005). However, there has been no systematic analysis of the effects of *Wolbachia* on the activity of different TEs in different host genotypes.

*Wolbachia* is a maternally inherited endosymbiont that is prevalent in invertebrates, including insects, arachnids, and nematodes (Werren et al. 2008; Kaur et al. 2021). Multiple studies have documented *Wolbachia* prevalence (Clark et al. 2005; Riegler et al. 2005; Weeks et al. 2007) and load (López-Madrigal et al. 2019; Liu et al. 2021) in natural and laboratory populations of *Drosophila* hosts. Associated to its mode of transmission, *Wolbachia* can have important effects on host reproduction, being responsible for phenomena such as cytoplasmic incompatibility, feminization, and male killing (Werren et al. 2008; Kaur et al. 2021). *Wolbachia* can also affect other aspects of host biology, including resistance to viral infection (Teixeira et al. 2008), gut microbiome composition (Simhadri et al. 2017), thermal preference (Truitt et al. 2019), sleep behaviour (Bi et al. 2018), and fecundity and lifespan (Serga et al. 2021). At the molecular level, *Wolbachia* is known to affect host gene expression (Baião et al. 2019; Biwot et al. 2020), and meiotic recombination rate (Singh, 2019), as well as the aforementioned TE-related properties Mayoral et al. 2014; Touret et al. 2014; Riegler et al. 2005).

Here, we test the impact of *Wolbachia* on TE expression by using host lines where *Wolbachia* is present versus where it was removed. Specifically, we use flies from the *Drosophila melanogaster* Genetic Reference Panel (DGRP), a panel of isogenic lines derived from a natural population, whose genomes have been fully sequenced and annotated for TE insertions (Mackay et al. 2012; Rahman et al. 2015). We selected 25 DGRP lines that were naturally infected with *Wolbachia* for which we estimated *Wolbachia* loads in individual flies and recorded the number of TE insertions for 14 TEs, representing different families. We found differences in *Wolbachia* loads and in number of novel TE insertions between genotypes, as well as an association between the two. We then generated a *Wolbachia*-free counterpart for each of the 25 target genotypes and used our panel of 25 paired *Wolbachia*-positive and *Wolbachia*-negative lines to quantify transcription levels of the 14 target TEs. We found variation in TE expression depending on host genotype, TE identity, and *Wolbachia* status. Whether *Wolbachia* removal led to increased or decreased TE expression appeared to be more of a property of host genotype than of TE identity.

## Results and Discussion

To investigate the effect of *Wolbachia* infection on TE activity, we focused on 25 *Drosophila melanogaster* genotypes, for which we documented differences in *Wolbachia*loads and in number of insertions of 14 target TEs (Fig. 1). We then generated a corresponding panel of 25 lines from which *Wolbachia* was cleared, and compared expression level of our target TEs in adult females between the pairs of *Wolbachia*- positive (Wolb+) and *Wolbachia*-negative (Wolb-) flies (Fig. 2).

**Fig. 1:**
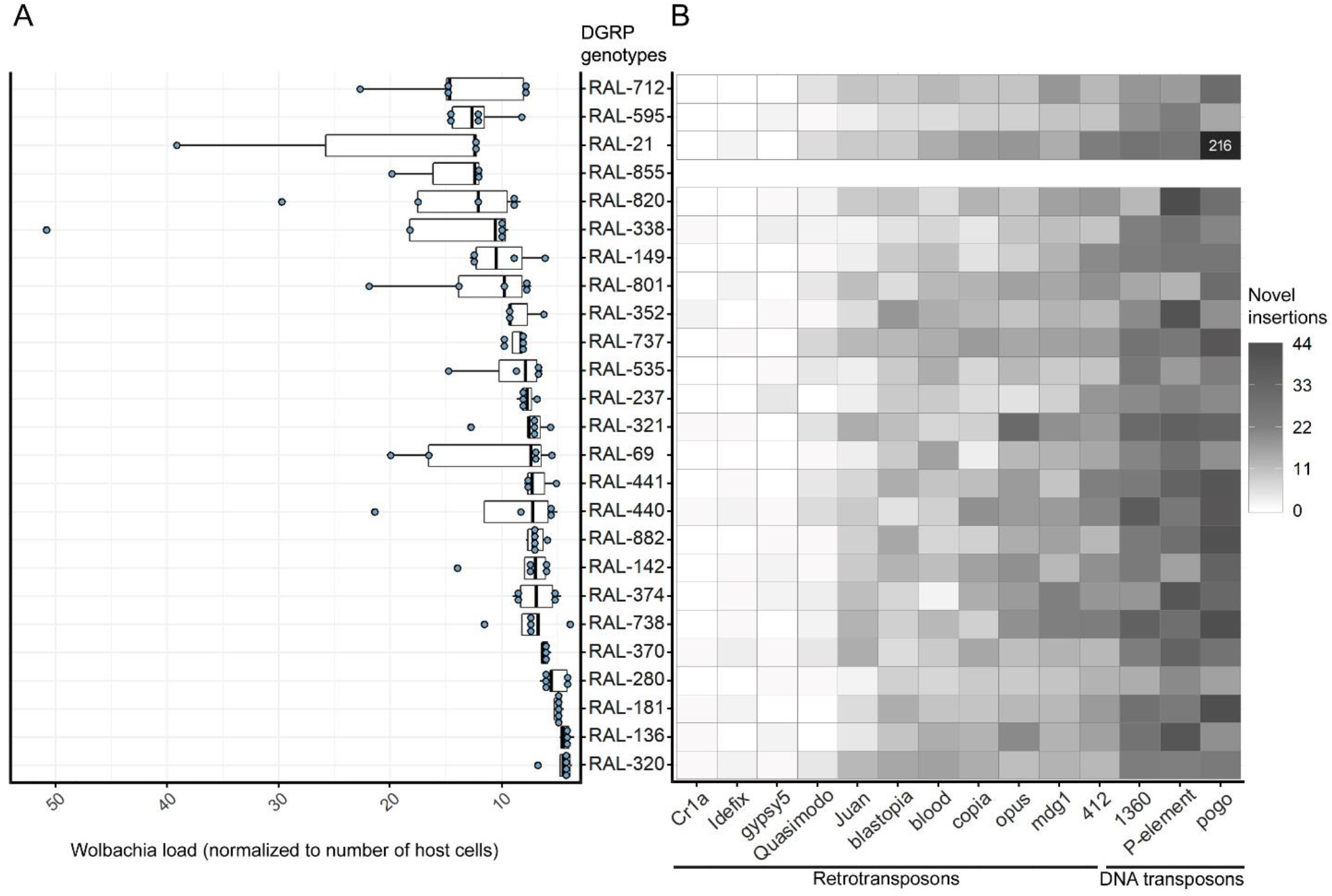
Characterization of our 25 target host lines in relation to *Wolbachia* load in individual flies (*A*) and to the number of novel TE insertions in their genomes (*B*). The 25 genotypes are organized along the y-axis in order of the median value of *Wolbachia* load. (*A*) *Wolbachia* load relative to number of host cells (x-axis). Each blue dot is a biological replicate and represents one single female. *Wolbachia* load is significantly different across genotypes (ANOVA; F_24,83_ =2.252, p-value=0.004). (*B*) Heatmap representing the predicted number of novel insertions for our 14 target TEs. TEs are organized in the x-axis by median number of novel insertions across genotypes. The scale of grey, from white to dark grey, represents, respectively, from the lowest to highest number of novel insertions. The TE pogo in genotype RAL-21 is out of the scale, with 216 novel insertions annotated. There was no information in Rahman et al. (2015) for RAL-855. The number of novel insertions differed significantly between genotypes (ANCOVA; F_22,4974_=1.3e+28, p<2e-16), TEs (F_13,4974_=1.7e+29, p<2e-16) and *Wolbachia* titers (F_1,4974_=2.2e+27, p<2e-16).

**Fig. 2:**
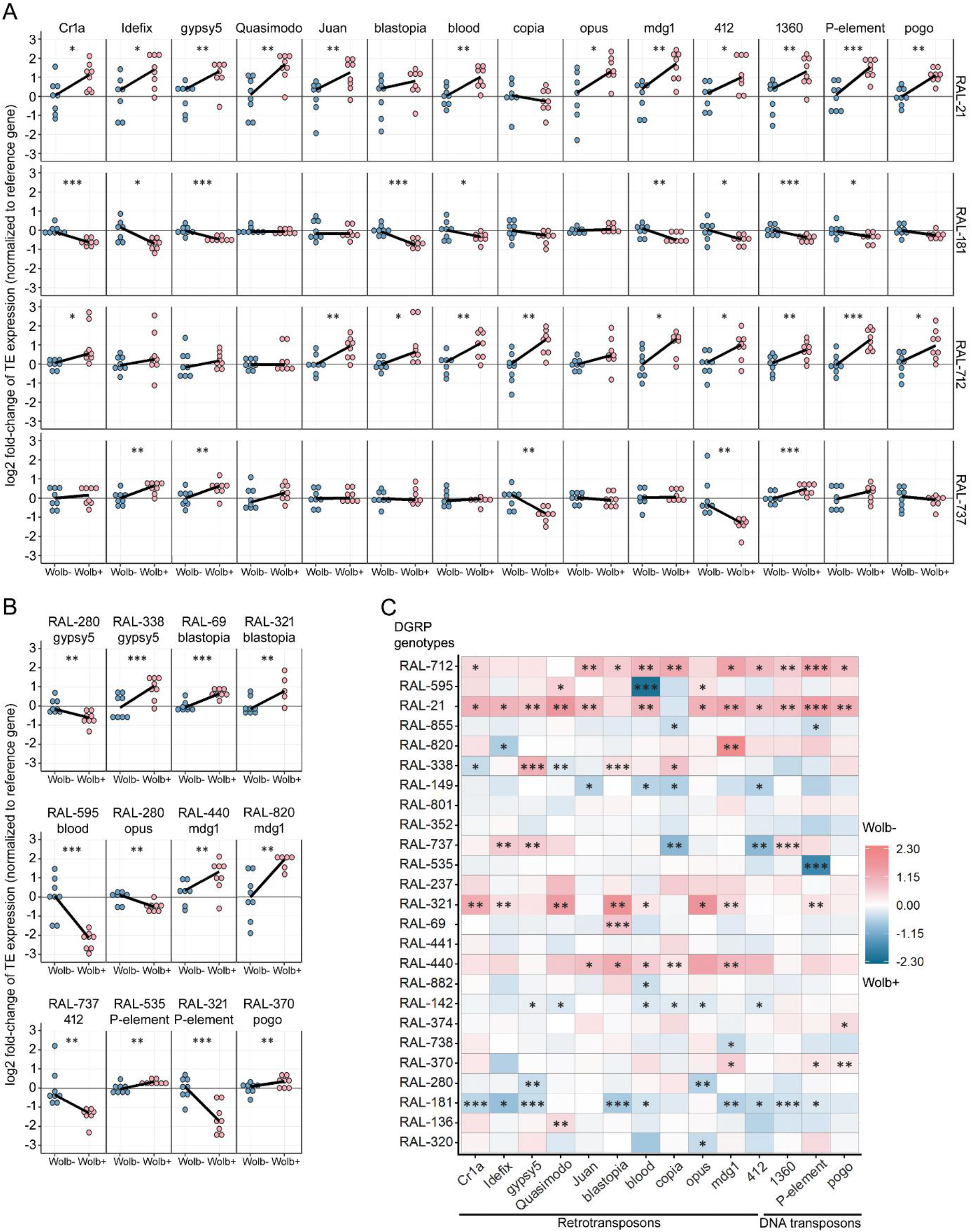
Expression levels of 14 TEs in adult female flies of 25 genotypes with (blue) versus without (pink) *Wolbachia*. TEs are ordered from left to right by median (or average, for tied medians) number of novel insertions. (*A*) Expression of the 14 TEs in genotypes RAL-21 and RAL-181. The same plots for all other genotypes can be found in Fig S2. (*B*) Expression of different TEs across various genotypes, illustrating cases where expression levels are statistically different between *Wolbachia* status. Each dot in plots (*A*) and (*B*) represents a biological replicate, corresponding to a pool of 10 female flies. Statistical significance for expression differences between Wolb+ and Wolb-is shown as * for p<0.05, ** for p<0.01, and *** for p<0.001 (Linear Mixed-Effects model, see Material and Methods). (*C*) Heatmap representing differences in expression level for the 14 target TEs between Wolb+ and Wolb-flies of all 25 different genotypes. Genotypes in the y-axis are ordered by *Wolbachia* load (as in Fig 1B). Cells are shown in a gradient from blue to pink, depending on whether expression was higher in Wolb+ relative to Wolb- (blue) or the other way around (pink).

### Host genotypes differ in Wolbachia loads and in number of TE insertions

We randomly selected 25 of the 85 DGRP lines known to be infected with the wMel strain of *Wolbachia* (Mackay et al. 2012; Richardson et al. 2012). For each of these lines, we measured *Wolbachia* loads in five individual adult females 10 days post-eclosion, the same sex and age used to measure TE expression. For this, we used quantitative realtime PCR (qPCR) with primers for one *Wolbachia-specific* gene (*wsp*), to estimate number of bacterial cells, and for one host-specific gene (*actin*), to assess number of host cells.

Across the ca. 125 flies assayed individually, *Wolbachia* loads varied between a minimum of 3.5 and a maximum of 51 *Wolbachia* cells per host cell. Only six individuals, of different genotypes, had greater than 20 *Wolbachia* per host cell. These estimates of *Wolbachia* density fall along the same order of magnitude as those found through sequencing of the DGRP lines (0.9 to 17.1 copies per host cell) (Richardson et al. 2012), or through qPCR of whole bodies (Bénard et al. 2021; Chrostek et al. 2021) and gonadal tissues (Correa and Ballard, 2012) of other *Drosophila melanogaster* genotypes, as well as for other *Wolbachia* strains (Chrostek and Teixeira, 2015).

We found differences in *Wolbachia* loads between host genotypes, with median values ranging from 5 to 15 copies of *Wolbachia* per host cell (Fig. 1A), and estimated broad-sense heritability (H^2^) of 0.22 (among-line variance=10.3, within-line variance=36.0). While little is known about what host loci harbour natural allelic variation contributing to variation in *Wolbachia* loads, we do know that loads vary with environmental factors, including temperature (Wiwatanaratanabutr and Kittayapong 2009), host diet (Ponton et al. 2015; Serbus et al. 2015), and viral infection (Kaur et al. 2020).

First, we validated *in silico* predictions of insertions (Mackay et al. 2012) by PCR with primers for the fly genomic sequence flanking 132 predicted novel insertions in 11 genotypes (Table S1). The amplicons from each of the insertions were sized (agarose gel) and sequenced to confirm the presence, length, and identity of the inserted DNA (Table S1). For 100% of the predicted insertion locations we tested, we confirmed the presence of a TE insertion, and, in most cases, we also confirmed that the size and the sequence of the inserted DNA corresponded to the predicted TE identity (Fig. S1 A). For 113 (85.6%) of the insertions tested, the inserted TE corresponded to the most likely expected identity (*cf*. the predictions made from the whole genome sequence data), and for 16, it corresponded to the second most likely TE (Mackay et al. 2012). We observed that 67 insertions (50.8%) had the size corresponding to the expected full length of that TE, 44 (33.3%) were smaller, and 21 (15.9%) were larger (Fig. S1A; Table S1).

With predictions of novel insertions validated, we used data from the TIDAL-FLY v1.0 tool of Rahman et al. (2015) to gather information about the number of novel insertions for each of our 14 target transposable elements in 24 of our 25 study genotypes (there were no data for genotype RAL-855). We found significant differences in the number of novel insertions between genotypes and *Wolbachia* titers. The retrotransposons *Cr1a, gypsy5*, and *Idefix* had the lowest number of predicted novel insertions (with zero for the majority of the lines), while the DNA transposons *1360, pogo*, and *P-element* generally had the highest number of novel insertions, in accordance with other studies describing DNA transposons as most active (Bourque et al. 2018). For most individual TEs, the estimated number of novel insertions varied between 0 and 44, with the exception of the TE pogo, predicted to have 216 novel insertions in the line RAL-21 (Fig. 1B; Rahman et al. 2015). Note that the *in silico* predictions of the number of TE insertions are likely to be underestimates of the actual number of insertions. The DGRP lines were originally sequenced using a combination of Illumina and 454 sequencing technologies (Mackay et al. 2012), which generate short-reads and, as such, are not ideal for detecting TE insertions (Panda and Slotkin 2020; Rech et al. 2022; Fiston-Lavier et al. 2015; Goerner-Potvin and Bourque 2018). Moreover, new insertions may also have occurred after sequencing. We confirmed experimentally multiple “false negatives” in TE insertion predictions for the DGRPs. By running PCRs with TE-specific primers and DNA from eight DGRP genotypes predicted in Mackay et al. (2012) to have no insertions (novel or shared) of particular TEs. In all 17 cases tested, we verified presence of those TEs (Fig. S1 B). However, even if predictions are underestimates of actual number of insertions, the effects should be similar/random across TEs and genotypes with equivalent sequence coverage depth.

### TE transcription level varies with *Wolbachia* status in a host genotype-dependent manner

Using qPCR with TE-specific primers and a reference host gene, we quantified the expression of our 14 target TEs in eight replicate pools of ten 10-days old females each, for each of the 25 Wolb+ and Wolb-pairs of genotypes (Cq data in Table S2). Expression levels differed significantly (Linear Mixed-Effects model) between TEs (F_13,4796_=4.75, p=2.97e-08), genotypes (F_24,4796_=26.63, p<2.2e-16), and with *Wolbachia* status (F_1,4796_=28.79, p=8.45e-08), with all interactions being significant (p<0.0001 in all cases) (Fig. 2).

Given the significant effect of *Wolbachia* status on TE expression, we then specifically compared expression of each of the 14 TEs in each of the 25 host genotypes with versus without *Wolbachia*. We found statistically significant differences in TE expression between Wolb+ and Wolb-lines for a total of 83 of the 350 (23,7%) genotype-TE combinations tested (Fig. 2; Fig. S2; Table S3). We observed distinct scenarios depending on TE and genotype: higher expression in Wolb-flies for 50 in 350 cases (14,3%) and higher expression in Wolb+ flies for 33 in 350 cases (9,4%).

For any given TE, the effect of *Wolbachia* on expression was not the same across genotypes, and, for any given genotype, the effect of *Wolbachia* on TE expression was not the same across TEs. However, for some genotypes, we observed some consistency in the effects of *Wolbachia* on TE expression. For genotypes RAL-149, RAL-142, and RAL-181, when statistically significantly different between Wolb+ and Wolb-flies, TE expression was always higher in Wolb- (blue shades in Fig. 2 C) relative to Wolb-flies. Conversely, for genotypes RAL-712, RAL-21, RAL-321, and RAL-440, when significantly different, TE expression was always higher in Wolb-(pink shades in Fig. 2C) relative to Wolb+flies. RAL-21 and RAL-712 stood out from the other genotypes analysed for having both naturally higher *Wolbachia* loads (Fig. 1 A) and higher levels of expression across most TEs after *Wolbachia* was removed (Fig. 2 A).

### Effect size data (when differences were statistically significant)

When significant, *Wolbachia* removal more often resulted in an increase than in a decrease in TE expression (pink shades versus blue shades in Fig. 2C, respectively), and effect sizes were generally stronger when expression was higher in Wolb-relative to Wolb+ flies. We found that when Wolbachia removal resulted in increased TE expression (pink shades in Fig 2C), genotypes almost doubled median expression levels of affected TEs (1.9 times higher expression in Wolb-relative to Wolb+; median log2 fold-change=1), going up to four times difference between medians for *mdg1* in RAL-820, and 19.7 times difference between individual samples for *opus* in RAL-321 (Fig. 2B and S2). Conversely, when Wolbachia removal resulted in a significant decrease in TE expression (blue shades in Fig. 2C), genotypes had a median of 1.4 times lower expression in Wolb-relative to Wolb+ (log2 fold-change=0.5), going down to 4.6 times difference between medians for *blood* in RAL-595, and 22.6 times (log2 fold-change=4.5) difference between individual samples for *412* in RAL-737 (Fig. 2B and S2).

## Conclusion

In this study, we investigated the hypothesis of a relationship between *Wolbachia* infection and TE activity. *Wolbachia* is a prevalent endosymbiotic bacterium whose impact on TE mobilization was suggested by distinct lines of evidence, including: 1) effect on piRNA expression (Mayoral et al. 2014), 2) effect on rate of transposition of retrotransposon gypsy (Touret et al. 2014), and 3) *Wolbachia* strain-replacement co-occurring with invasion of DNA transposon P-element (Riegler et al. 2005). We tested whether the expression of 14 diverse TEs was different between *Wolbachia-infected* and *Wolbachia*-free *D. melanogaster* flies of 25 distinct genotypes, which differed in *Wolbachia* loads and in number of TE insertions. We found significant differences in levels of TE transcript between Wolb+ and Wolb-flies in 23.7% of the 350 genotype-TE combinations analysed, amounting to a TE expression difference of 20 times for individual samples and 4 times for medians of same-genotype replicates. The effects of *Wolbachia* removal observed were not uniform for any given TE nor for most genotypes. However, some genotypes did stand out in having multiple TEs for which the direction *Wolbachia* effect on expression was the same.

As if often done (e.g., Becking et al. 2020; Torres et al. 2021), we used transcript level as a proxy for TE activity, reasoning that higher expression creates more opportunities for new insertions. However, transcription is only a necessary first step in TE mobilization, and it is conceivable that *Wolbachia*, or other factors, might impact TE integration post-transcription. Expression level was quantified using qPCR and following MIQE guidelines (Taylor et al. 2010), including technical and biological replication, ensuring template quality, careful selection of reference genes, and correction for differences in primer efficiency. Moreover, all samples being directly compared were ran together and using the same batch of reagents.

This study was performed using a subset of the DGRP lines, a panel of isogenic and fully sequenced genotypes that provide the possibility of looking at genotypic variation. Even though the genotypes are not naturally occurring, in that they were highly isogenised post-collection of a natural population, they represent naturally segregating allelic variants. Many studies have showed differences between DGRP genotypes for different traits, such as longevity (Ivanov et al. 2015), fecundity (Durlam et al. 2014), body size and thermal plasticity therein (Lafuente et al. 2018) resistance to viral (Magwire et al. 2012) and bacterial (Howick and Lazzaro 2017) infection, and sensitivity to oxidative stress (Weber et al. 2012), amongst other traits (Mackay and Huang 2018). Our results, showing differences between genotypes in Wolbachia loads and TE insertion number, as well as in the effect of *Wolbachia* removal on TE activity emphasize the importance of analysing multiple genotypes to have a more complete understanding of biological phenomena.

Novel genetic variants created by TE mobilization can be, and are often, deleterious (McFaddenaf et al. 1997; Hedges et al. 2007; Belancio et al. 2008; Ayarpadikannan and Kim 2014). As such, high TE activity can put natural populations under stable conditions at a disadvantage. On the other hand, TE insertions can also be beneficial and, particularly in conditions of environmental perturbation, TE activity could contribute to novel genetic variants better adjusted to the changed conditions (e.g., Rey et al. 2016). The question of which and how intrinsic and extrinsic factors affect TE activity is a fundamentally interesting and largely unresolved question, especially for animal when comparing to plant TEs (Thieme et al. 2017). It is also a question drawing considerable attention, and where we can expect to continue to gather valuable insights.

## Materials and Methods

### Confirming *in silico* predictions of TE insertions in DGRP lines

We looked to validate *in-silico* predictions both in terms of potential false positives (focusing on specific insertions) and potential false negatives (focusing on particular TEs deemed as having no insertions in some genotypes.

First, we selected 132 of the predicted novel insertions in 12 of the DGRP lines and designed primers for the sequence flanking those insertions (Table S1). For each of the lines, we extracted gDNA from pools of 10 males (homogenised using pestles), using DNeasy Blood & Tissue kit (Qiagen), following manufacturer’s instructions. We then used 4ng of this gDNA in 15μl Long PCR reactions with 0.5μM primers, 2% DMSO, 0.5mM dNTPs mix, 0.21μL of GoTaq enzyme (Promega). Thermocycler conditions included 2 min at 92°C; 10 cycles of 92°C for 10 sec, 60°C for 15 sec, 68°C for 10 min; 30 cycles of 92°C for 15 sec, 60°C for 30 sec, 68°C for 10 min + 20 sec cycle elongation for each successive cycle; 7 min at 68°C. Amplicons were sized (1% agarose gel electrophoresis) and sequenced (ThermoFisher BigDye Terminator v1.1, or SUPREMErun^™^ from; same forward primers used for amplification) and these were NZYTech compared with the size and sequence of the canonical *Drosophila* transposons (Flybase version 9.42).

Second, we tested the absence of specific TEs in genotypes annotated as having no insertions of that TE. We ran PCR with primers specific for each of seven TEs (*blood*, *copia, gypsy5, H-element, jockey, opus, pogo;* Table S4) and gDNA extracted from pools of ten adult females (extractions as described above) of eight genotypes (RAL-109, RAL-161, RAL-237, RAL-350, RAL-362, RAL-555, RAL-776, RAL-808) predicted to not have one or more of those TEs (Mackay et al. 2012), confirming presence or absence of insertion band in 1% agarose gel (Fig. S1B), With the gDNA from each of the target genotypes, we ran two types of positive controls: 1) with TE-specific primers with gDNA extracted from a genotype (RAL-321) predicted to have insertions of all six TEs, and 2) with primers for the *Drosophila* gene *RPL32* (Table S4) present in every line. gDNA extracted as described above was used in 10μL PCR reactions containing 0.4ng gDNA, 0.25U GoTaq (Promega), 1.5mM MgCl_2_, and 0.5μM of each primer. The thermal cycling protocol was: 10 min at 95°C; 35 cycles of 95°C for 30 sec, 60°C for 1 min, 72°C for 30 sec; 5 min at 72°C.

### Fly lines and husbandry

For each of the lines selected, we generated a *Wolbachia*-free version following procedures described in Teixeira et al. (2008) and Chrostek et al. (2013). In short, flies were first rid of *Wolbachia* by feeding on food supplemented with tetracycline antibiotic (0.05mg/mL) for two generations. Their gut flora was then restored by placing sterilized eggs of *Wolbachia*-cleared flies (10 min in 50% bleach followed by washing in sterilized water) on food supplemented with a bacterial inoculum (150μL of a mix prepared by mixing 2mL of sterile water with 1g of a month-old food filtered to remove eggs and larvae) of each respective untreated (*Wolbachia*-positive) line. Flies were *Wolbachia*- free and gut microbiota-homogenized for at least five generations before the experiments were initiated.

Flies were reared at 25°C and 12h:12h light:dark cycle, in vials with cornmeal-agar food (45g/L molasses, 75g/L white sugar, 70g/L corn flour, 20g/L yeast extract, 10g/L agar-agar and 25mL Nipagin at 10%) and similar density conditions. For our experiments, we transferred newly eclosed adult flies to vials in groups of 10 females and 6 males. Females were sampled for extraction of DNA (for quantification of *Wolbachia*) or of RNA (for quantification of TE expression) at 10 days of age.

### *Wolbachia* presence and loads

We used *Wolbachia*-specific primers against the *Wolbachia surface protein* gene (*wsp*; sequence from Teixeira et al. 2008) to confirm that the tetracycline-treated Wolb-lines were indeed *Wolbachia*-free and to quantify *Wolbachia* loads in the untreated Wolb+ lines.

We confirmed the absence of *Wolbachia* in each of the tetracycline-treated lines in 10μL PCR reactions, containing 0.4ng gDNA template, 0.25U GoTaq (Promega), 1.5mM MgCl_2_, and 0.5μM of each primer (*wsp*). We used gDNA extracted (QIAGEN’s DNeasy Blood & Tissue Kit, following manufacturer’s indications) from three pools of ten females (mixed ages) from each of the 25 Wolb-lines, homogenized using QIAGEN Tissue Lyser II (2 min at 23s/f). As positive control, we extracted gDNA from the 25 Wolb+ lines (same protocol) and used those samples as template. Thermal cycle was 4 min at 95°C; 35 cycles of 95°C for 30 sec, 60°C for 1 min and 72°C for 30 sec; 5 min at 72C^°^. PCR amplicons were checked by electrophoresis gel (1% agarose) and we confirmed the successful removal of *Wolbachia* (no amplicon) in all 25 target DGRP lines.

We measured *Wolbachia* loads in five individual females (10 days post-eclosion) from each of our 25 target Wolb+ lines. Individual females were homogenized in QIAGEN ATL Buffer in 96 well-plates with a sterile glass bead per well on a Tissue Lyser II (QIAGEN) at 23s/f for 2 minutes, before DNA was extracted using the Quick-DNA^TM^ 96 Kit (Zymo Research), following manufacturer’s instructions. DNA was eluted in 200μL Buffer AE from the kit and stored at −20oC until qPCR, which was run using primers (Table S4) for either a *Wolbachia* gene (*wsp*; measuring *Wolbachia* load) or for a host gene (*actin*, proxy for number of host cells) as described below.

### RNA extraction and cDNA synthesis for TE expression quantification by qPCR

To quantify TE expression, we extracted RNA from eight replicate pools of 10 co-housed, 10-day old females, for each of the 25 pairs of Wolb+ and Wolb-genotypes (total of 50 lines). Whole bodies were homogenized in 400 μL TRIzol (Invitrogen) using a sterile glass bead in microcentrifuge tubes and a Tissue Lyser II (Qiagen) at 26s/f for 1 minute. Homogenates were stored at −80°C until further processing. Once thawed, we added 80 μL of chloroform, centrifuged (12.000g for 15 min at 4°C) and collected the supernatant aqueous phase containing the RNA (to avoid carrying fly tissues and fat to the RNA extraction step), and then we added 400 μL more TRIzol. Total RNA was then extracted using the Direct-zol^TM^ 96 RNA Kit (Zymo Research), following manufacturer instructions. We used 4μg of RNA to synthesize cDNA with NZY First-Strand cDNA Synthesis Kit (NZYTech), following manufacturer’s instructions. cDNA was then diluted 1:10 in sterile water (Sigma) to be used as template in qPCR with primers (Table S4) against each of the 14 target TEs (*412*, *1360, blastopia, blood, copia, Cr1a, gypsy5, Idefix, Juan, mdg1, opus, Quasimodo, P-element, pogo*) or against one reference gene, *EF1*, chosen from a number of candidates (*18S*, *Act5c, actin, EF1, ELF2, Gapdh1, Mnf, Rpl32, Rps20, TBP, tubulin*) using Normfinder (Andersen et al. 2004) and selecting a gene with Cq values similar to that of the TEs being tested (qPCR reagents and thermocycle as described below).

### qPCR with standard curves

We used qPCR to measure both *Wolbachia* titers (gDNA template and primers for one *Wolbachia*-specific gene, *wsp*, and one host-specific reference gene, *actin*) and TE expression (cDNA template and primers for each of the 14 target TEs and one reference gene). Template preparation and primers were described above.

For each biological replicate sample, we ran two technical replicate reactions in an QuantStudioTM 7 Flex Real-Time PCR System (Applied Biosystems™). We used 4μL of genomic template, 0.5μL of each primer (0.2μM) and 5μL of SYBR Green I^®^ (Bio Rad), and the following thermal cycling conditions: 2 min at 50°C; 10 min at 95°C; 40 cycles of 95°C for 30 sec, 60°C for 1 min and 72°C for 30 sec. We discarded biological replicates for which the standard deviation between Cq values of the two technical replicates was greater than 0.5, and calculated the mean Cq value between technical replicates for each of all other biological replicates. Processing of Cq data of biological replicates is detailed below.

For each gene and each TE, we also obtained standard curves relating amount of template and Cq values. These were obtained by using as template a 1:10 serial dilution (8 dilutions) of a cleaned (Macherey-Nagel’s NucleoSpin Gel and PCR Clean-up) and quantified (Invitrogen’s Qubit™) PCR product (obtained by PCR on gDNA extracted from flies from the standard line Oregon R). We used the equations for the linear regression of log quantity of starting template (x-axis) and Cq value (y-axis) to: 1) do absolute quantification of *wsp* and *actin*, as there in no obvious calibrator sample for analysis of *Wolbachia* loads, and 2) calculate primer efficiency required for the relative quantification of TE expression using the Pfaffl method (2001).

### Processing qPCR Cq data to quantify *Wolbachia* and TE expression

For the absolute quantification of *Wolbachia* loads, we used the mean Cq values of each biological sample and the standard curves for *wsp* and actin to determine the quantity of each of the genes in the sample used as template: quantity =10^((Cq-b)/m)^, where b is the intercept and m is the slope of the linear regression equation. We estimated the quantity of both *wsp* and *actin* in each sample and then calculated the ratio between the two (quantity of *wsp* / quantity of *actin*) as a measurement of *Wolbachia* load in relation to host cells.

For the relative quantification of TE expression, we used the Pfaffl method: expression ratio = E_(TE)_^ΔCq(TE)^/ E_(EF1)_^ΔCq(EF1)^. E is the amplification efficiency for each primer pair and is calculated based on the equation of the linear regression of the respective standard curve: E=10^-1/slope^ (primer efficiencies in Table S4). For all TE x genotype samples, ΔCq refers to the difference in Cq values between a calibrator sample (average of same-genotype Wolb+ samples) and each sample for that genotype (Wolb- and Wolb+).

### Statistical analysis

Statistical analyses were performed in R (version 3.6.3), using Rstudio (version 1.4.1106).

To test for differences in *Wolbachia* loads across the 25 target genotypes we used ANOVA with genotype as fixed factor and replicate as random factor: *aov*(*Wolbachia* load ~ genotype + (1|replicate)) in R syntax. We estimated broad sense heritability (H^2^) for *Wolbachia* loads as H^2^=σ^2^A/(σ^2^A +σ^2^W), in which σ^2^A is the among-line variance and σ^2^W is the within-line variance. Variance components were extracted using the *VCA R* package (Schuetzenmeister and Dufey 2020). To test for differences in the number of novel insertions we used ANCOVA with *Wolbachia* load as covariate and genotype and TE as fixed factor: *aov*(novel insertions ~ mean *Wolbachia* load + genotype * TE) in R syntax.

To account for variation in TE expression with *Wolbachia* status, we used ANCOVA with TE, genotype, and *Wolbachia* status as fixed factors: *aov*(log2(expression ratio) ~ TE * genotype * *Wolbachia* status) in R syntax. Then, for each TE and genotype, we compared expression between Wolb+ and Wolb-flies using the Linear Mixed-Effects model (packages *lme4*, Bates et al. 2015, and *lmerTest*, Kuznetsova et al. 2017) with *Wolbachia* status as fixed factor and biological replicate as random factor: *lmer*(log2(expression ratio) ~ *Wolbachia* status + (1|replicate)) in R syntax.

## Supporting information

Table S2 - available upon acceptance

## Acknowledgements

We want to acknowledge Renato Alves for helping with bioinformatic tools, Daniel Sobral for aligning next generation sequencing reads against TE canonical sequences, the Fly Facility at Instituto Gulbenkian de Ciência for fly maintenance, and Élio Sucena and Isabel Gordo for important feedback.

## Funding

This work was supported by the Portuguese science funding agency, Fundação para a Ciência e Tecnologia (FCT): PhD fellowships for AT Eugénio (SFRH/BD/115535/2016, COVID/BD/151707/2021), PhD fellowships for MSP Marialva (SFRH/BD/51882/2012), and research grants to P Beldade (PTDC/BIA-EVF/0017/2014, PTDC/BIA-EVL/0321/2021).

## Notes

### Competing Interest Statement

The authors have declared no competing interest.

## References

1. Andersen CL, Jensen JL, Orntoft TF. 2004. Normalization of real-time quantitative reverse transcription-PCR data: A model-based variance estimation approach to identify genes suited for normalization, applied to bladder and colon cancer data sets. Cancer Res 64:5245–5250.

2. Anderson SN, Stitzer MC, Zhou P, Ross-Ibarra J, Hirsch CD, Springer NM. 2019. Dynamic patterns of transcript abundance of transposable element families in maize. G3-Genes Genom Genet 9:3673–3682.

3. Aravin AA, Hannon GJ, Brennecke J. 2007. The Piwi-piRNA pathway provides an adaptive defense in the transposon arms race. Science 318:761–764.

4. Ayarpadikannan S, Kim HS. 2014. The impact of transposable elements in genome evolution and genetic instability and their implications in various diseases. Genomics Inform 12:98–104.

5. Baião GC, Schneider DI, Miller WJ, Klasson L. 2019. The effect of *Wolbachia* on gene expression in *Drosophila paulistorum* and its implications for symbiont-induced host speciation. BMC Genomics 20:465.

6. Bates D, Maechler M, Bolker B, Walker S. 2015. Fitting linear mixed-effects models using lme4. J Stat Softw 67:1–48.

7. Becking T, Gilbert C, Cordaux R. 2020. Impact of transposable elements on genome size variation between two closely related crustacean species. Annal Biochem 600: 113770.

8. Belancio VP, Hedges DJ, Deininger P. 2008. Mammalian non-LTR retrotransposons: for better or worse, in sickness and in health. Genome Res 18:343–358.

9. Bénard A, Henri H, Noûs C, Vavre F, Kremer N. 2021. *Wolbachia* load variation in *Drosophila* is more likely caused by drift than by host genetic factors. Peer Community Journal 1:e38.

10. Bennetzen JL, Wang H. 2014. The contributions of transposable elements to the structure, function, and evolution of plant genomes. Annu Rev Plant Biol 65:505–530.

11. Bi J, Sehgal A, Williams JA, Wang YF. 2018. *Wolbachia* affects sleep behavior in *Drosophila melanogaster*. J Insect Physiol 107:81–88.

12. Biwot JC, Zhang HB, Liu C, Qiao JX, Yu XQ, Wang YF. 2020. *Wolbachia-induced* expression of kenny gene in testes affects male fertility in *Drosophila melanogaster*. J Insect Sci 27:869–882.

13. Bourque G, Burns KH, Gehring M, Gorbunova V, Seluanov A, Hammell M, Imbeault M, Izsvák Z, Levin HL, Macfarlan TS et al. 2018. Ten things you should know about transposable elements. Genome Biol 19:199.

14. Chen JE, Cui G, Wang X, Liew YJ, Aranda M. 2018. Recent expansion of heat-activated retrotransposons in the coral symbiont *Symbiodinium microadriaticum*. ISME J 12:639–643.

15. Choi J, Lyons DB, Kim MY, Moore JD, Zilberman D. 2020. DNA methylation and histone H1 jointly repress transposable elements and aberrant intragenic transcripts. Mol Cell 77:310–323.e7.

16. Chrostek E, Marialva MSP, Esteves SS, Weinert LA, Martinez J, Jiggins FM, Teixeira L. 2013. *Wolbachia* variants induce differential protection to viruses in *Drosophila melanogaster*: A phenotypic and phylogenomic analysis. PLoS Genet 9:e1003896.

17. Chrostek E, Martins N, Marialva MS, Teixeira L. 2021. *Wolbachia-conferred* antiviral protection is determined by developmental temperature. mBio 12:e0292320.

18. Chrostek E, Teixeira L. 2015. Mutualism breakdown by amplification of *Wolbachia* genes. PLoS Biol 13:e1002065.

19. Clark ME, Anderson CL, Cande J, Karr TL. 2005. Widespread prevalence of *Wolbachia* in laboratory stocks and the implications for *Drosophila* research. Genetics 170:1667–1675.

20. Correa CC, J. Ballard WO. 2012. *Wolbachia* gonadal density in female and male *Drosophila* vary with laboratory adaptation and respond differently to physiological and environmental challenges. J Invertebr Pathol 111:197–204.

21. Durham M, Magwire M, Stone E. Leips J. 2014. Genome-wide analysis in *Drosophila* reveals age-specific effects of SNPs on fitness traits. Nat Commun 5:4338.

22. Emera DM, Wagner GP. 2012. Transposable element recruitments in the mammalian placenta: impacts and mechanisms. Brief Funct Genomics 11:267–276.

23. Santos M, Braasch I, Boileau N, Meyer BS, Sauteur L, Böhne A, Belting HG, Affolter M, Salzburger W. 2014. The evolution of cichlid fish egg-spots is linked with a cis-regulatory change. Nat Commun. 5:5149.

24. Fiston-Lavier AS, Barrón MG, Petrov DA, González J. 2015. T-lex2: genotyping, frequency estimation and re-annotation of transposable elements using single or pooled next-generation sequencing data. Nucleic Acids Res 43:27.

25. Goerner-Potvin P, Bourque G. 2018. Computational tools to unmask transposable elements. Nat Rev Genet 19:688–704.

26. González J, Karasov TL, Messer PW, Petrov DA. 2010. Genome-wide patterns of adaptation to temperate environments associated with transposable elements in *Drosophila*. PLoS Genet 6:33–35.

27. González J, Petrov DA. 2009. The adaptive role of transposable elements in the *Drosophila* genome. Gene 448:124–133.

28. Goodier JL. 2016. Restricting retrotransposons: a review. Mob DNA 7:16.

29. Guio L, Barrón MG, González J. 2014. The transposable element Bari-Jheh mediates oxidative stress response in *Drosophila*. Mol Ecol 23:2020–2030.

30. Guio L, González J. 2019. New insights on the evolution of genome content: population dynamics of transposable elements in flies and humans. In: Anisimova M, editors. Evolutionary Genomics. Methods in Molecular Biology. Humana, New York. 1910:505–530.

31. Guzzardo PM, Muerdter F, Hannon GJ. 2013. The piRNA pathway in flies: highlights and future directions. Curr Opin Genet Dev 23:44–52.

32. Habibi L, Shokrgozar MA, Tabrizi M, Modarressi MH, Akrami SM. 2014. Mercury specifically induces LINE-1 activity in a human neuroblastoma cell line. Mutat Res Genet Toxicol Environ Mutagen 759:9–20.

33. Hedges DJ, Deininger PL. 2007. Inviting instability: Transposable elements, double-strand breaks, and the maintenance of genome integrity. Mutat Res 616:46–59.

34. Howick VM, Lazzaro BP. 2017. The genetic architecture of defence as resistance to and tolerance of bacterial infection in *Drosophila melanogaster*. Mol Ecol 26:1533–1546.

35. Ivanov DK, Escott-Price V, Ziehm M, Magwire MM, Mackay TF, Partridge L, Thornton JM. 2015. Longevity GWAS using the *Drosophila* genetic reference panel. J Gerontol A Biol Sci Med Sci 70:1470–1478.

36. Kaur RJ, Shropshire D, Cross KL, Leigh B, Mansueto AJ, Stewart V, Bordenstein SR, Bordenstein SR. 2021. Living in the endosymbiotic world of *Wolbachia*: A centennial review. Cell Host Microbe 29:879–893.

37. Kaur R, Martinez J, Rota-Stabelli O, Jiggins FM, Miller WJ. 2020. Age, tissue, genotype and virus infection regulate *Wolbachia* levels in *Drosophila*. Mol Ecol 29:2063–2079.

38. Kuznetsova A, Brockhoff PB, Christensen RHB. 2017. lmerTest package: tests in linear mixed effects models. J Stat Softw 82:1–26.

39. Lafuente E, Duneau D, Beldade P. 2018. Genetic basis of thermal plasticity variation in *Drosophila melanogaster* body size. PLoS Genet 14:e1007686.

40. Le Thomas A, Rogers AK, Webster A, Marinov GK, Liao SE, Perkins EM, Hur JK, Aravin AA, Tóth KF. 2013. Piwi induces piRNA-guided transcriptional silencing and establishment of a repressive chromatin state. Genes & Dev 27:390–399.

41. Liu XC, Li ZX. 2021. Transmission of the wMel *Wolbachia* strain is modulated by its titre and by immune genes in *Drosophila melanogaster* (*Wolbachia* density and transmission). J Invertebr Pathol 181:107591.

42. López-Madrigal S, Duarte EH. 2019. Titer regulation in arthropod-*Wolbachia* symbioses. FEMS Microbiol Lett 366:fnz232.

43. Mackay TFC, Huang W. 2018. Charting the genotype–phenotype map: lessons from the *Drosophila melanogaster* Genetic Reference Panel. Wiley Interdiscip Rev Dev Biol 7:e289.

44. Mackay TFC, Richards S, Stone EA, Barbadilla A, Avroles JF, Zhu D, Casillas S, Han Y, Magwire MM, Cridland JM et al. 2012. The *Drosophila melanogaster* genetic reference panel. Nature 482:173–178.

45. Magwire MM, Fabian DK, Schweyen H, Cao C, Longdon B, Bayer F, Jiggins FM. 2012. Genome-wide association studies reveal a simple genetic basis of resistance to naturally coevolving viruses in *Drosophila melanogaster*. PLoS Genet 8:e1003057.

46. Mayoral JG, Etebari K, Hussain M, Khromykh AA, Asgari S. 2014. *Wolbachia* infection modifies the profile, shuttling and structure of MicroRNAs in a mosquito cell line. PLoS One 9:e96107.

47. McFaddenaf J, Knowlesb G. 1997. Escape from evolutionary stasis by transposon-mediated deleterious mutations. J Theor Biol 186:441–447.

48. Mérel V, Boulesteix M, Fablet M, Vieira C. 2020. Transposable elements in *Drosophila*. Mob DNA 11:23.

49. Miousse IR, Chalbot MCG, Lumen A, Ferguson A, Kavouras IG, Koturbash I. 2015. Response of transposable elements to environmental stressors. Mutat Res Rev Mutat Res 765:19–39.

50. Newman MR, Sykes PJ, Blyth BJ, Bezak E, Lawrence MD, Morel KL, Ormsby RJ. 2014. The methylation of DNA repeat elements is sex-dependent and temporally different in response to X radiation in radiosensitive and radioresistant mouse strains. Radiat Res 181:65–75.

51. Panda K, Slotkin RK. 2020. Long-read cDNA sequencing enables a “gene-like” transcript annotation of transposable elements. Plant Cell 32:2687–2698.

52. Pereira JF, Ryan PR. 2019. The role of transposable elements in the evolution of aluminium resistance in plants. J Exp Bot 70:41–54.

53. Petrov DA, Schutzman JL, Hartl DL, Lozovskaya ER. 1995. Diverse transposable elements are mobilized in hybrid dysgenesis in *Drosophila virilis*. PNAS 92:8050–8054.

54. Pfaffl MW. 2001. A new mathematical model for relative quantification in real-time RT-PCR. Nucleic Acids Res 29:e45.

55. Ponton F, Wilson K, Holmes A, Raubenheimer D, Robinson KL, Simpson SJ. 2015. Macronutrients mediate the functional relationship between *Drosophila* and *Wolbachia*. Proc R Soc 282:2014–2029.

56. Rahman R, Chirn G, Kanodia A, Sytnikova YA, Brembs B, Bergman CM, Lau NC. 2015. Unique transposon landscapes are pervasive across *Drosophila melanogaster* genomes. Nucleic Acids Res 43:10655–10672.

57. Rech GE, Radío S, Guirao-Rico S, Aguilera L, Horvath V, Green L, Lindstadt H, Jamilloux V, Quesneville H, González J. 2022. Population-scale long-read sequencing uncovers transposable elements associated with gene expression variation and adaptive signatures in *Drosophila*. Nat Commun 13:1948.

58. Rep M, van der Does HC, Cornelissen BJC. 2005. Drifter, a novel, low copy hAT-like transposon in *Fusarium oxysporum* is activated during starvation. Fungal Genet Biol 42:546–553.

59. Rey O, Danchin E, Mirouze M, Loot C, Blanchet S. 2016. Adaptation to global change: A transposable element–epigenetics perspective. Trends Ecol Evol 31:514–526.

60. Richardson MF, Weinert LA, Welch JJ, Linheiro RS, Magwire MM, Jiggins FM, Bergman CCM. 2012. Population genomics of the *Wolbachia* endosymbiont in *Drosophila melanogaster*. PLoS Genet 8:e1003129.

61. Riegler M, Sidhu M, Miller WJ, O’Neill SL. 2005. Evidence for a global *Wolbachia* replacement in *Drosophila melanogaster*. Curr Biol 15:1428–1433.

62. Schuetzenmeister A, Florian Dufey F. 2020. VCA: Variance Component Analysis. R package version 1.4.3.

63. Serbus LR, White PM, Silva JP, Rabe A, Teixeira L, Albertson R, Sullivan W. 2015. The impact of host diet on *Wolbachia* titer in *Drosophila*. PLoS Pathog 11:e1004777.

64. Serga SV, Maistrenko OM, Matiytsiv NP, Vaiserman AM, Kozeretska IA. 2021. Effects of *Wolbachia* infection on fitness-related traits in *Drosophila melanogaster*. Symbiosis 83:163–172.

65. Serrato-Capuchina A, Matute DR. 2018. The role of transposable elements in speciation. Genes 9:254.

66. Sienski G, Dönertas D, Brennecke J. 2012. Transcriptional silencing of transposons by Piwi and maelstrom and its impact on chromatin state and gene expression. Cell 151:964–980.

67. Signor S. 2020. Transposable elements in individual genotypes of *Drosophila simulans*. Ecol Evol 10:3402–3412.

68. Simhadri RK, Fast EM, Guo R, Schultz MJ, Vaisman N, Ortiz L, Bybee J, Slatko BE, Frydman HM. 2017. The gut commensal microbiome of *Drosophila melanogaster* is modified by the endosymbiont *Wolbachia*. mSphere 2:e00287–17.

69. Taylor S, Wakem M, Dijkman G, Alsarraj M, Nguyen M. 2010. A practical approach to RT-qPCR – Publishing data that conform to the MIQE guidelines. Methods 50:S1–S5.

70. Teixeira L, Ferreira Á, Ashburner M. 2008. The bacterial symbiont *Wolbachia* induces resistance to RNA viral infections in *Drosophila melanogaster*. PLoS Biol 6:e1000002.

71. Thieme M, Lanciano S, Balzergue S, Daccord N, Mirouze M, Bucher E. 2017. Inhibition of RNA polymerase II allows controlled mobilisation of retrotransposons for plant breeding. Genome Biol 18:134.

72. Torres DE, Thomma BPHJ, Seidl MF. 2021. Transposable elements contribute to genome dynamics and gene expression variation in the fungal plant pathogen *Verticillium dahlia*. GBE 13:evab135.

73. Tóth KF, Pezic D, Stuwe E, Webster A. 2016. The piRNA pathway guards the germline genome against transposable elements. In: Wilhelm D, Bernard P, editors. Non-coding RNA and the reproductive system. Advances in Experimental Medicine and Biology. Springer, Dordrecht. 886.

74. Touret F, Guiguen F, Terzian C. 2014. *Wolbachia* influences the maternal transmission of the gypsy endogenous retrovirus in *Drosophila melanogaster*. MBio 5:e01529–14.

75. Trizzino M, Park Y, Holsbach-Beltrame M, Aracena K, Mika K, Caliskan M, Perry GH, Vincent J. Lynch VJ, Brown CD. 2017. Transposable elements are the primary source of novelty in primate gene regulation. Genome Res 27:1623–1633.

76. Truitt AM, Kapun M, Kaur R, Miller WJ. 2019. *Wolbachia* modifies thermal preference in *Drosophila melanogaster*. Environ Microbiol 21:3259–3268.

77. van’t Hof AE, Campagne P, Rigden DJ, Yung CJ, Lingley J, Quail MA, Hall N, Darby AC, Saccheri IJ. 2016. The industrial melanism mutation in British peppered moths is a transposable element. Nature 534:102–105.

78. Venner S, Feschotte C, Biémont C. 2009. Dynamics of transposable elements: towards a community ecology of the genome. Trends Genet 25:317–323.

79. Weber AL, Khan GF, Magwire MM, Tabor CL, Mackay TFC, Anholt RRH. 2012. Genome-wide association analysis of oxidative stress resistance in *Drosophila melanogaster*. PLoS One 7:e34745.

80. Weeks AR, Turelli M, Harcombe WR, Reynolds KT, Hoffmann AA. 2007. From parasite to mutualist: Rapid evolution of *Wolbachia* in natural populations of *Drosophila*. PLoS Biol 5:e114.

81. Werren JH, Baldo L, Clark ME. 2008. *Wolbachia:* master manipulators of invertebrate biology. Nat Rev Microbiol 6:741–751.

82. Wiwatanaratanabutr I, Kittayapong P. 2009. Effects of crowding and temperature on *Wolbachia* infection density among life cycle stages of *Aedes albopictus*. J Invertebr Pathol 102:220–224.

