## Supplementary Figures for "Effects of *Wolbachia* on transposable element activity largely depend on *Drosophila melanogaster* host genotype"

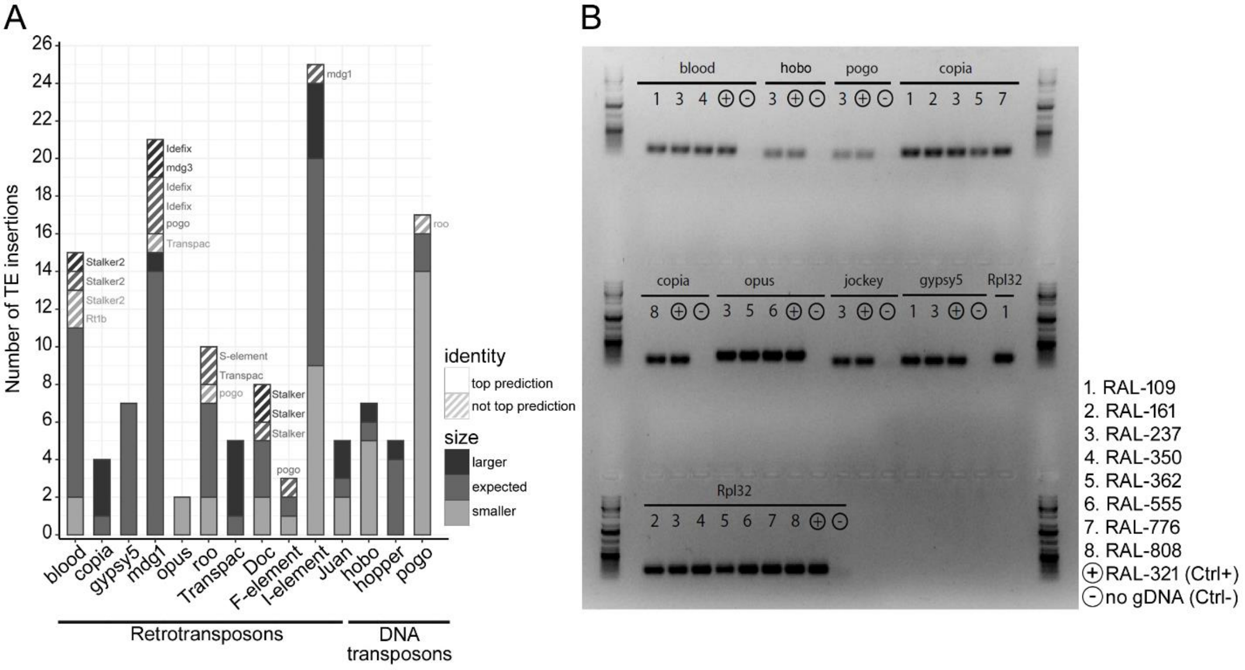


**Fig. S1**: Confirmation of *in silico* predictions of TE insertions in the DGRP lines (Mackay et al. 2012). (*A*) Confirmation of the identity and size of 132 predicted TE insertions. For each TE, the histogram represents the number of insertions we did and did not confirm the identity to be the most likely cf. the *in silico* prediction (solid versus pattern fill), and whether the size of the inserted fragment corresponded to that of the predicted TE (shade of grey). For the cases where the inserted TE did not correspond to the top prediction (pattern fill), the figure includes the name of the actual TE identity confirmed by sequencing the inserted fragment. (*B*) Electrophoresis gel of PCR amplicons confirming the presence (amplification with TE specific primers) of insertions in DGRPs (numbers) described as not having specific TEs (names on top of gel lanes). The positive control (Ctrl+) corresponds to PCR using as template DNA from a line predicted to contain insertions of all the TEs studied (RAL-321), and the negative control (Ctrl-) corresponds to PCR run with primers for each TE but without gDNA. For each DNA sample, we also ran PCR using primers for a host gene present in all lines (Rpl32).

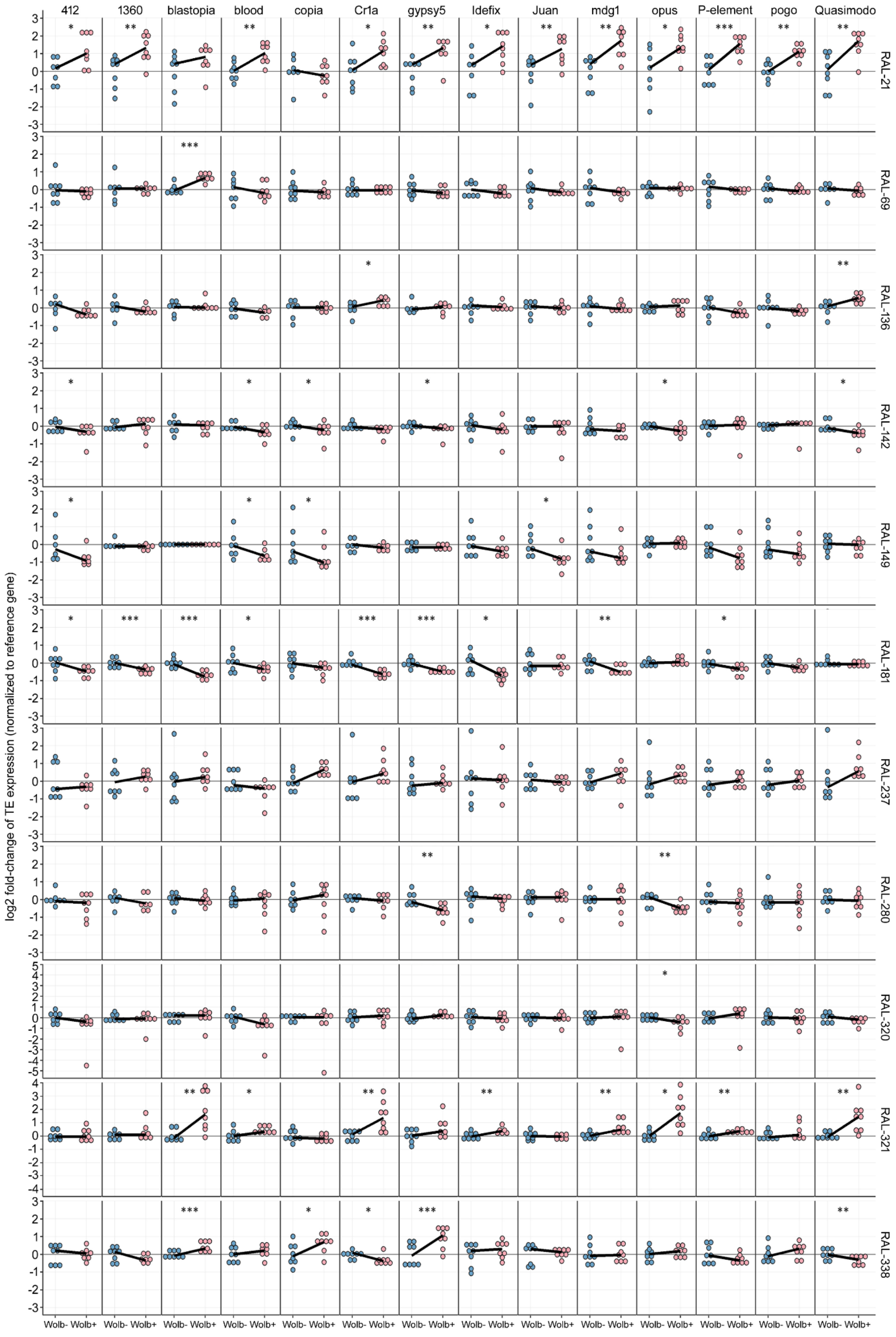
412
1360
blastopia
blood
copia
Cr1a
gypsy5
Idefix
Juan
mdg1
opus
P-element
pogo
Quasimodo
RAL-21
RAL-69
RAL-136
RAL-142
RAL-149
RAL-181
RAL-237
RAL-280
RAL-320
RAL-321
RAL-338
log2 fold-change of TE expression (normalized to reference gene)
Wolb- Wolb+

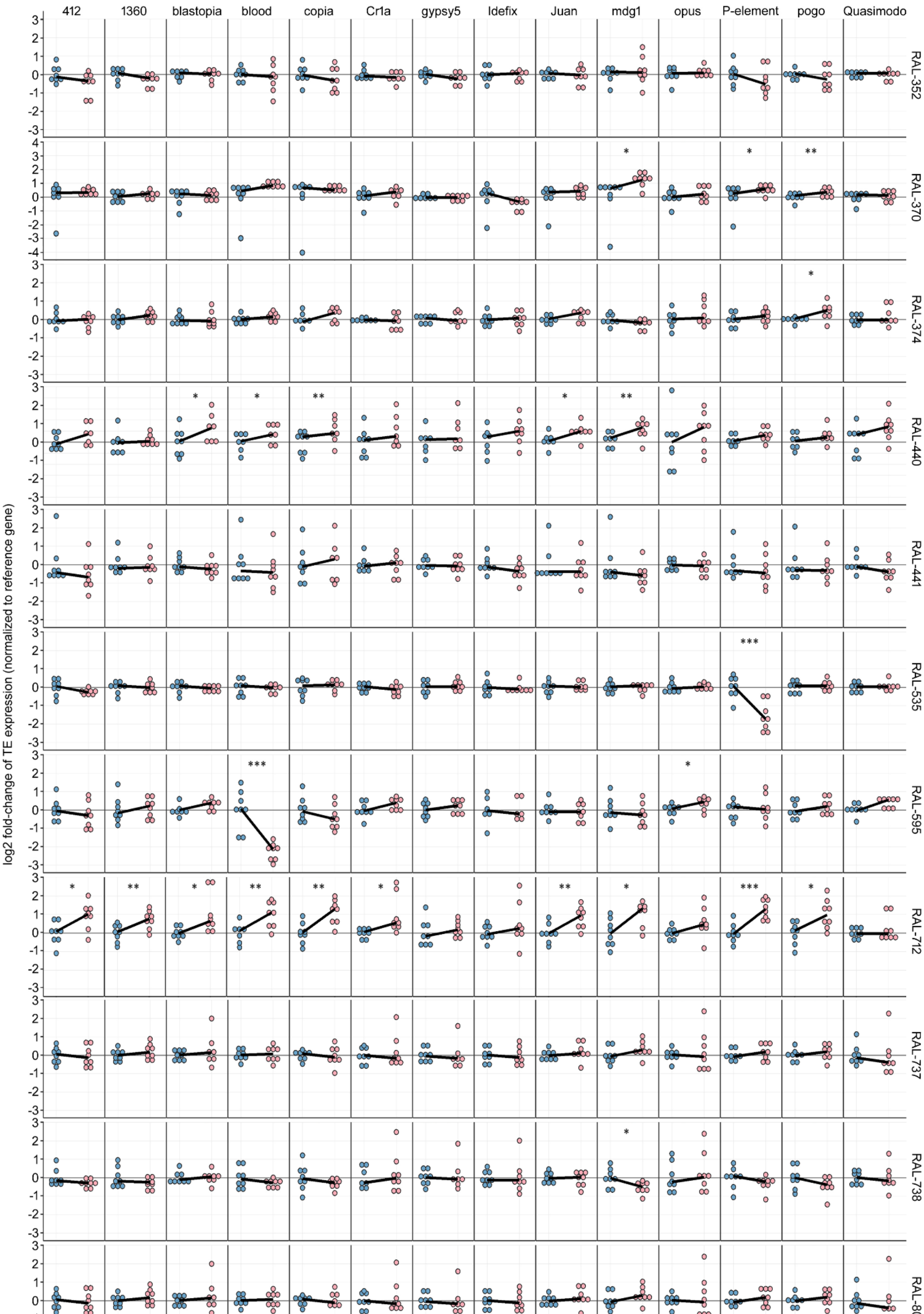

412
1360
blastopia
blood
copia
Cr1a
gypsy5
Idefix
Juan
mdg1
opus
P-element
pogo
Quasimodo
RAL-352
RAL-370
RAL-374
RAL-440
RAL-441
RAL-535
RAL-595
RAL-712
RAL-737
RAL-738
RAL-801
log2 fold-change of TE expression (normalized to reference gene)
Wolb- Wolb+

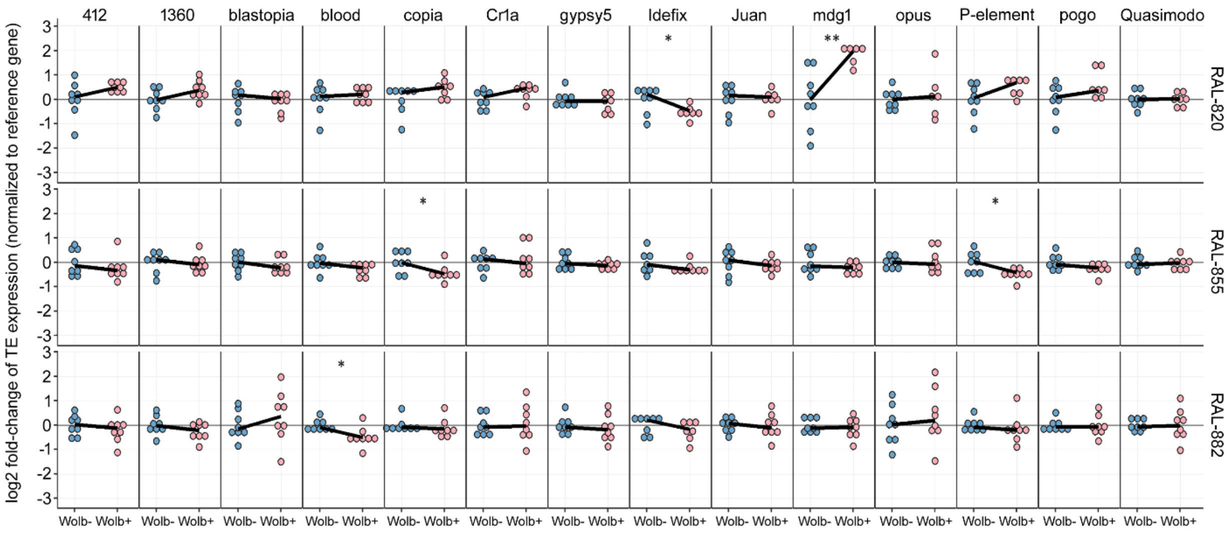


**Fig. S2**: Complete panel of expression levels for the 14 target TEs in adult female flies from our 25 target lines with (blue; Wolb-) versus without (pink; Wolb-) Wolbachia. Each panel corresponds to data for one TE (labels on top) in one genotype (labels to the right). Each dot in a panel represents a biological replicate, corresponding to a pool of 10 female flies, and black lines join medians for Wolb+ and Wolb- samples. Statistical significance for expression differences between Wolb+ and Wolb- is shown as * for p<0.05, ** for p<0.01, and *** for p<0.001 (Linear Mixed-Effects model, see Material and Methods; actual t and p-values in Table S3). Note that, for completeness, this figure includes panels also represented in Fig. 2A and 2B.
