## Supplementary material for "Effects of *Wolbachia* on transposable element activity largely depend on *Drosophila melanogaster* host genotype": Table S1

**Supplementary Table S1**

**Experimental confirmation of predicted TE insertions**

| Nr | TE insertion position | DGRP | Identity predicted (Mackay et al. 2012) | Identity confirmation (sequencing) |
|---|---|---|---|---|
| 1 | 2L:14056691..14056894 | RAL-321 | *blood* | no (Stalker2) |
| 2 | 2R:4932328..4932331 | RAL-321 | *blood* | yes |
| 3 | 3L:1030460..1030467 | RAL-321 | *blood* | yes |
| 4 | 3L:22887401..22887413 | RAL-321 | *blood* | no (Rt1b) |
| 5 | 3R:7387389..7387392 | RAL-358 | *blood* | no (Stalker2) |
| 6 | 2L:9310321..9310326 | RAL-790 | *blood* | yes |
| 7 | 2L:18683411..18683415 | RAL-790 | *blood* | yes |
| 8 | 2R:20107115..20107118 | RAL-790 | *blood* | yes |
| 9 | 3L:5167028..5167031 | RAL-790 | *blood* | yes |
| 10 | 3R:7994453..7994456 | RAL-790 | *blood* | yes |
| 11 | 3R:19792545..19792549 | RAL-790 | *blood* | yes |
| 12 | 3R:22107018..22107022 | RAL-790 | *blood* | yes |
| 13 | X:10922194..10922200 | RAL-790 | *blood* | no (Stalker2) |
| 14 | 3R:4991260..4991264 | RAL-810 | *copia* | yes |
| 15 | 3R:4991260..4991264 | RAL-812 | *copia* | yes |
| 16 | 3R:13161944..13161944 | RAL-908 | *copia* | yes |
| 17 | 2L:4426305..4426310 | RAL-908 | *copia* | yes |
| 18 | 2L:19806748..19807008 | RAL-790 | *gypsy5* | yes |
| 19 | 2R:15116726..15116809 | RAL-790 | *gypsy5* | yes |
| 20 | 3L:773447..773450 | RAL-790 | *gypsy5* | yes |
| 21 | 3L:3891219..3891222 | RAL-790 | *gypsy5* | yes |
| 22 | 3L:9736885..9736886 | RAL-790 | *gypsy5* | yes |
| 23 | 3R:2368919..2368922 | RAL-790 | *gypsy5* | yes |
| 24 | 3R:22031220..22031223 | RAL-790 | *gypsy5* | yes |
| 25 | 2L:1788665..1788852 | RAL-357 | *mdg1* | no (Idefix) |
| 26 | 2L:15102151..15102178 | RAL-357 | *mdg1* | yes |
| 27 | 2R:19003049..19003053 | RAL-357 | *mdg1* | yes |
| 28 | 2R:19347971..19347974 | RAL-357 | *mdg1* | no (mdg3) |
| 29 | 3L:10095528..10095532 | RAL-357 | *mdg1* | yes |
| 30 | 3L:20245853..20245856 | RAL-357 | *mdg1* | yes |
| 31 | 3R:5798180..5798183 | RAL-357 | *mdg1* | yes |
| 32 | 3R:12117132..12117136 | RAL-357 | *mdg1* | yes |
| 33 | 3R:23192134..23192145 | RAL-357 | *mdg1* | no (Idefix) |
| 34 | X:345626..345630 | RAL-357 | *mdg1* | no (Transpac) |
| 35 | X:16472629..16472643 | RAL-357 | *mdg1* | yes |
| 36 | 2L:19305699..19305703 | RAL-908 | *mdg1* | yes |
| 37 | 2R:11793310..11793699 | RAL-908 | *mdg1* | yes |
| 38 | 3L:676970..677090 | RAL-908 | *mdg1* | yes |
| 39 | 3L:13514785..13514789 | RAL-908 | *mdg1* | yes |
| 40 | 3R:5737883..5737886 | RAL-908 | *mdg1* | yes |
| 41 | 3R:8620803..8620806 | RAL-908 | *mdg1* | yes |
| 42 | 3R:16918669..16918673 | RAL-908 | *mdg1* | yes |
| 43 | 3R:18045826..18045829 | RAL-908 | *mdg1* | yes |
| 44 | X:22997736..22997762 | RAL-908 | *mdg1* | no (Idefix) |
| 45 | X:5024909..5024914 | RAL-908 | *mdg1* | no (pogo) |
| 46 | 2L:7336324..7336327 | RAL-908 | *opus* | yes |
| 47 | 2R:17950637..17950706 | RAL-443 | *opus* | yes |
| 48 | 3R:17957511..17957520 | RAL-358 | *roo* | yes |
| 49 | 3R:7884988..7884992 | RAL-358 | *roo* | yes |
| 50 | 3R:17515331..17515334 | RAL-358 | *roo* | no (Transpac) |

| | | | | |
|---|---|---|---|---|
| 51 | X:21270475..21270477 | RAL-908 | *roo* | no (pogo) |
| 52 | 2L:9204672..9204675 | RAL-908 | *roo* | yes |
| 53 | 2L:9361897..9361901 | RAL-908 | *roo* | yes |
| 54 | 2L:9705174..9705181 | RAL-908 | *roo* | yes |
| 55 | 2L:12191481..12191488 | RAL-908 | *roo* | no (S-element) |
| 56 | 2L:12434248..12434266 | RAL-908 | *roo* | yes |
| 57 | 2L:12902268..12902274 | RAL-908 | *roo* | yes |
| 58 | 2R:12108723..12108729 | RAL-810 | *Transpac* | yes |
| 59 | 3L:12856836..12856842 | RAL-908 | *Transpac* | yes |
| 60 | X:345626..345630 | RAL-357 | *Transpac* | yes |
| 61 | X:345626..345630 | RAL-892 | *Transpac* | yes |
| 62 | X:15333499..15333504 | RAL-443 | *Transpac* | yes |
| 63 | 2L:11138677..11138734 | RAL-357 | *Doc* | yes |
| 64 | 2L:11138677..11138734 | RAL-892 | *Doc* | yes |
| 65 | 3R:7873179..7873180 | RAL-357 | *Doc* | no (Stalker) |
| 66 | 3R:7873179..7873180 | RAL-381 | *Doc* | no (Stalker) |
| 67 | 3R:7873179..7873180 | RAL-761 | *Doc* | no (Stalker) |
| 68 | 2R:7936918..7936946 | RAL-810 | *Doc* | yes |
| 69 | 3R:10301789..10301822 | RAL-908 | *Doc* | yes |
| 70 | 2L:1399467..1399478 | RAL-908 | *Doc* | yes |
| 71 | 3R:6947766..6947992 | RAL-810 | *F-element* | no (pogo) |
| 72 | 3L:7996067..7996071 | RAL-908 | *F-element* | yes |
| 73 | 3R:9994118..9994126 | RAL-908 | *F-element* | yes |
| 74 | 2R:10573276..10573286 | RAL-357 | *I-element* | yes |
| 75 | 2R:13861586..13861600 | RAL-810 | *I-element* | yes |
| 76 | 3L:5068011..5068084 | RAL-908 | *I-element* | yes |
| 77 | 3R:11405981..11405987 | RAL-357 | *I-element* | yes |
| 78 | 3R:14271683..14271693 | RAL-810 | *I-element* | yes |
| 79 | 2L:11388784..11388793 | RAL-790 | *I-element* | yes |
| 80 | 2R:7961110..7961122 | RAL-790 | *I-element* | yes |
| 81 | 2R:12732718..12732730 | RAL-790 | *I-element* | yes |
| 82 | 2R:14371197..14371209 | RAL-790 | *I-element* | yes |
| 83 | 3R:4277178..4277189 | RAL-790 | *I-element* | yes |
| 84 | 3R:14671496..14671508 | RAL-790 | *I-element* | yes |
| 85 | 3R:21118597..21118611 | RAL-790 | *I-element* | yes |
| 86 | 3R:21756397..21756400 | RAL-790 | *I-element* | yes |
| 87 | 2R:2648988..2649001 | RAL-908 | *I-element* | yes |
| 88 | 2R:7224003..7224015 | RAL-908 | *I-element* | yes |
| 89 | 3L:4501669..4501675 | RAL-908 | *I-element* | no (mdg1) |
| 90 | 3L:5068011..5068084 | RAL-908 | *I-element* | yes |
| 91 | 3L:10088380..10088381 | RAL-908 | *I-element* | yes |
| 92 | 3L:22408430..22408442 | RAL-908 | *I-element* | yes |
| 93 | 3R:8071617..8071628 | RAL-908 | *I-element* | yes |
| 94 | 3R:11192794..11192806 | RAL-908 | *I-element* | yes |
| 95 | 3R:22596610..22596622 | RAL-908 | *I-element* | yes |
| 96 | 3R:22850421..22850434 | RAL-908 | *I-element* | yes |
| 97 | X:302449..302464 | RAL-908 | *I-element* | yes |
| 98 | X:10027285..10027298 | RAL-908 | *I-element* | yes |
| 99 | 2R:5572279..5572291 | RAL-810 | *Juan* | yes |
| 100 | 3L:17698435..17698491 | RAL-357 | *Juan* | yes |
| 101 | 3L:17698435..17698491 | RAL-761 | *Juan* | yes |
| 102 | X:18924423..18924436 | RAL-908 | *Juan* | yes |
| 103 | 3L:16130441..16130456 | RAL-443 | *Juan* | yes |
| 104 | 3R:10703373..10703379 | RAL-357 | *hobo* | yes |
| 105 | 3R:22133514..22133521 | RAL-357 | *hobo* | yes |

| | | | | |
|---|---|---|---|---|
| 106 | 2R:12871101..12871113 | RAL-810 | *hobo* | yes |
| 107 | 3L:19555349..19555354 | RAL-761 | *hobo* | yes |
| 108 | 3L:19555349..19555354 | RAL-810 | *hobo* | yes |
| 109 | 3L:19555349..19555354 | RAL-812 | *hobo* | yes |
| 110 | 3R:22134885..22134892 | RAL-908 | *hobo* | yes |
| 111 | 2L:2390528..2390533 | RAL-357 | *hopper* | yes |
| 112 | 3R:13938897..13938901 | RAL-357 | *hopper* | yes |
| 113 | 3R:13938897..13938901 | RAL-812 | *hopper* | yes |
| 114 | 3R:15010776..15010780 | RAL-908 | *hopper* | yes |
| 115 | X:4110151..4110442 | RAL-810 | *hopper* | yes |
| 116 | 2R:11767856..11767857 | RAL-810 | *pogo* | yes |
| 117 | 2R:11767856..11767857 | RAL-381 | *pogo* | yes |
| 118 | 3L:21388576..21388578 | RAL-810 | *pogo* | yes |
| 119 | 3L:21388576..21388578 | RAL-812 | *pogo* | yes |
| 120 | 3R:11292096..11292097 | RAL-357 | *pogo* | yes |
| 121 | 3R:2926760..2926761 | RAL-357 | *pogo* | yes |
| 122 | X:6015256..6015259 | RAL-908 | *pogo* | yes |
| 123 | X:6015256..6015259 | RAL-381 | *pogo* | yes |
| 124 | 3L:1571691..1571694 | RAL-358 | *pogo* | yes |
| 125 | 2L:12470302..12470309 | RAL-908 | *pogo* | yes |
| 126 | 2L:13770337..13770338 | RAL-908 | *pogo* | yes |
| 127 | 2L:13780514..13780515 | RAL-908 | *pogo* | yes |
| 128 | 2L:15151373..15151377 | RAL-908 | *pogo* | yes |
| 129 | 2L:18860089..18860091 | RAL-908 | *pogo* | yes |
| 130 | 2R:1749442..1749443 | RAL-908 | *pogo* | yes |
| 131 | 2R:5741587..5741594 | RAL-908 | *pogo* | yes |
| 132 | 2R:6426442..6426443 | RAL-908 | *pogo* | no (roo) |

| Supplementary Table S1 | | | | |
|---|---|---|---|---|
| **Experimental confirmation of predicted TE insertions** | | | | |
| **Nr** | **Size predicted (bp)** | **Size confirmation (Long-PCR; bp)** | **Forward primer (5' – 3')** | **Reverse primer (5' – 3')** |
| 1 | 7410 | yes (7000) | CGGGGATGTTGTCTTTCGG | TGCAATTCATCACCCCAAGT |
| 2 | 7410 | yes (7000) | CCTTGTCCATGAACACGGG | ACAACACTGCGGTTTCCA |
| 3 | 7410 | no, smaller (700) | TAAGTTTACCGCCAGCACG | GCGACACTTTTCCTCCTGT |
| 4 | 7410 | no, smaller (4000) | AGTAAATAGTCGGGGCGGG | CGATGGCAATCCTCCATTTG |
| 5 | 7410 | no, smaller (4000) | TCCAGCGGGCTCCAATTTA | CCGTCCCAGACTATATCACC |
| 6 | 7410 | yes (7000) | CAGCTCTCATGATGACGACG | GACCGAAATACACCGTTCCA |
| 7 | 7410 | yes (7000) | GACACCAGTACTTGACGCA | CAATGACCTGGCATCGCT |
| 8 | 7410 | yes (7000) | GACTCGAGTTCTTTACGCCA | AAGGACAAAGGATGCGGAT |
| 9 | 7410 | yes (7000) | TGTCGCATAAATCCACTCCG | CCCCATAATTCCCCGCAAT |
| 10 | 7410 | yes (7000) | GGTCAGAGCAAACCGGAAAA | CGTGTCGGCCCAGAATAT |
| 11 | 7410 | yes (7000) | ATCTGCTCTGGGGCCTCA | GGCCGTTGAGACTGAATTG |
| 12 | 7410 | no, smaller (1000) | AGTCAGGGCATGTGGTACTA | CTAAAAGGACCTTCTGCCCC |
| 13 | 7410 | no, higher (10000) | AGCATGCTTCAAAGGCAGG | AAGTGGAAGCAAGGAGTCC |
| 14 | 5100 | no, higher (6000) | TCCTCTCCCCCTCTCTGTCT | TTAAGCCCAACCACATAGCC |
| 15 | 5100 | no, higher (6000) | TCCTCTCCCCCTCTCTGTCT | TTAAGCCCAACCACATAGCC |
| 16 | 5100 | no, higher (6000) | CACGTGTCCATAGCCCATTT | CTGCTTAACCATTGCGTCCT |
| 17 | 5100 | yes (5000) | CTCGAGAGTTCGGAAAGCAT | AGGACTCTGGACAGGTGGTG |
| 18 | 7369 | yes (7000) | GCGTCTTAGCCATGATGAAGAA | TTGCCAAGTTTACGCTACG |
| 19 | 7369 | yes (7000) | GTTTTGTGTGATCGCGCTG | AGTGGGAGTTGAAGGGGTG |
| 20 | 7369 | yes (7000) | TGAACGTGAACTGAGAGTGG | CGGGCTCGCTGAATTGTC |
| 21 | 7369 | yes (7000) | GCTGAAAGAGGAGCCCAG | GGCAAGCTGGTGTATTGAAA |
| 22 | 7369 | yes (7000) | TTGGATGGGTAGGCAGTAGA | GCGTGGGAGTTATTGCGTA |
| 23 | 7369 | yes (7000) | TTTGCCGTTAGAAGCTGGAG | GCCACTGGACCCCACTTA |
| 24 | 7369 | yes (7000) | GCTAATACCGCCAGCCTTT | GATGGAGCTGGCCTTGAAA |
| 25 | 7480 | yes (7000) | TGACTTTCTTTGTGCGCCC | TCTCCTTAAACGTGGCTAGTG |
| 26 | 7480 | yes (7000) | GACCACCGTCGACTTTAAAA | GAGGAGCCGAGATCTGTC |
| 27 | 7480 | yes (7000) | TACGGCCTCCCCAAGTGAA | GGATATGACTGGTGCTTCTTATC |
| 28 | 7480 | no, higher (10000) | ACTTGGCCTAGGGAGCAATA | AGTTCAGTCCCAGAGGCC |
| 29 | 7480 | yes (7000) | AGAGGCCTGTGAGATCCTTT | AAGTGTTGCCCCATTCGAA |
| 30 | 7480 | yes (7000) | TGCCCGTACATTTATGCAGC | CGTGTGTGTGGAGCGATC |
| 31 | 7480 | yes (7000) | AGGGCACACAGTTTTGGTAC | CTCGAATGGCAACCTGTACT |
| 32 | 7480 | yes (7000) | CCAGAACGTCGACCAACC | GTGACCTAACGCAATACACAA |
| 33 | 7480 | no, higher (10000) | CCTCAACAAGTCGGCGGG | CCCCATTCCAACGCACAA |
| 34 | 7480 | no, smaller (5000) | TACAAGTTGGGGCACAGGA | TCGCAGCGGGAATATGAATT |
| 35 | 7480 | yes (7000) | TAAAGGCGAGAGGAAAGAGC | TTTTGCCCCGTTCGAGTG |
| 36 | 7480 | yes (7000) | AACTCCACCGACTTGCTTTT | TGGCTCTGACATGACCTAGA |
| 37 | 7480 | yes (7000) | AAAGGAGCAGGAGAAGGAGT | GAGGCAAAACAACGAGCATC |
| 38 | 7480 | yes (7000) | ATATTCACGGTTTTGGCCAG | GAGGGGCCGAGTCCTAAAT |
| 39 | 7480 | yes (7000) | CGCGCTTTGTTAACGTGG | CATATTGGCTCGCGTTTTGT |
| 40 | 7480 | yes (7000) | GAACACGCCGAGGATAGC | CGAAAATCTACTCAGCCTGCT |
| 41 | 7480 | yes (7000) | CTGACTGAGAAATGGGGCG | TCACCAAAGTTGCAGTCGAA |
| 42 | 7480 | yes (7000) | ACCGGGTGCCAAGGAATG | CTTCCTCTTCGCTTCAGTGG |
| 43 | 7480 | no, higher (10000) | ACTCTACGGCAAATGAATGGG | GGCGTGACCCTTACTGAA |
| 44 | 7480 | yes (7000) | AAAAGAGTCCGCAAGTCCAG | AGCTGTGTAACCGGGTAAGA |
| 45 | 7480 | yes (7000) | ACAGAGAGCGACGAAAGC | CGATGCGATGTCATGTCC |
| 46 | 7500 | no, smaller (700) | GCATGACGATTACGTGGCTA | ACAACCAAACGCTTTTCACC |
| 47 | 7500 | no, smaller (3000) | ATATGTCCTCGCCTGACCTG | GTTTCCACTGCACAGCCATA |
| 48 | 9092 | no, smaller (700) | ATCAGCGTTTAGGTCGATCG | CGATGATGCCCAGTCTTAAAT |
| 49 | 9092 | yes (10000) | CCCACATAGTTGCATGCCA | GTGTGTGGCAGTGGTAGT |
| 50 | 9092 | yes (10000) | CAGTGCCCCATTGTTACGA | CGAACTCATTGCAACTCCCT |

| | | | | |
|---|---|---|---|---|
| 51 | 9092 | no, smaller (1500) | TCGTTCTCTCAGCAACCAAC | AGAGAACCGAAACGCTACTG |
| 52 | 9092 | no, smaller (7000) | TTGGCGGGGAACAAATACAC | TGCCCCTTACCGCTCCAT |
| 53 | 9092 | yes (10000) | TCTTGTTTCCCCGCATCATG | TCTACCTCCGGAGCTCACA |
| 54 | 9092 | yes (10000) | CAACTGATGGGGTAAGCAATG | GCGAGCAAAGACAGCACTA |
| 55 | 9092 | yes (10000) | CGGTGCTTGAGGTGTCCC | CACTTCAAGCCTGGTTACCC |
| 56 | 9092 | yes (10000) | CACGACGTTAGAGCTGGAAC | CACGCATCAAGTCAGGGTT |
| 57 | 9092 | yes (10000) | GAATTTCGGGCTGGTTTTCA | CCATGATGTCCTGTGCTCC |
| 58 | 5200 | no, higher (6000) | ATAAACCGCAGCAAAAGTGG | CCAATGGATTTTCGAGAGGA |
| 59 | 5200 | no, higher (6000) | CCCACTTCCTCTTCCACTCA | AGTCGACCAGGGACAATGAC |
| 60 | 5200 | yes, (5000) | TAACGATGGTGGCTGCTACA | AAGGAAAGCGATTCAAGACC |
| 61 | 5200 | no, higher (6000) | TAACGATGGTGGCTGCTACA | AAGGAAAGCGATTCAAGACC |
| 62 | 5200 | no, higher (6000) | CTGCAACTTTCCATGGCTTT | ACAGCTTTCCCCTTCTGGAT |
| 63 | 4700 | no, smaller (3000) | AAAATCCATTCGGCAAACTG | TCGATCAGCGCCTAGTATCA |
| 64 | 4700 | yes (5000) | AAAATCCATTCGGCAAACTG | TCGATCAGCGCCTAGTATCA |
| 65 | 4700 | no, higher (7000) | ATTGTCTGCGCAACTGTCTG | ATGAATTCGTCTGCCTGTCC |
| 66 | 4700 | no, higher (7000) | ATTGTCTGCGCAACTGTCTG | ATGAATTCGTCTGCCTGTCC |
| 67 | 4700 | yes (5000) | AGCTTTAGCCACAGCCACAT | GAGAGGCACGCAGGTAAGAC |
| 68 | 4700 | yes (5000) | CGAAGACATCAGTCCTGCAA | CCGCTGACTGTGATTGCTAA |
| 69 | 4700 | no, smaller (3000) | GCACGAGACTCACACAGGAA | TTATGGCCATTGTACGCTGA |
| 70 | 4700 | yes, (5000) | TGCATCTGTGTGCGTATGTG | GCACTTTTTGCCTCTGTTCC |
| 71 | 4700 | yes (5000) | TAGGCGCTGTTATTGAAACC | CAGTGAAAGTGGGTGCAAA |
| 72 | 4700 | no, smaller (3500) | GGGATTTGCTCTTGCTCTTG | GCCATGGTCGAAACAAAACT |
| 73 | 4700 | yes (5000) | GCTTGTCAAAGGGTCCAAGA | TGTTATGTGCGCGAACTTGT |
| 74 | 5300 | no, smaller (700) | TCCGTCGGCTCTTATTTGTC | CGTCTTACACTCGCAGCAAA |
| 75 | 5300 | no, smaller (1500) | CCCAGATTCGCAATACCAAA | AACAAAAGCAACCACCAAGG |
| 76 | 5300 | no, smaller (700) | TCGAATTGATACAACCCCAAT | CTACTACGGCGGTGTTGGTT |
| 77 | 5300 | yes (5000) | GGCAGTGCAAACAAAAACAA | CTGAGGCCAAGGACTTATGC |
| 78 | 5300 | no, smaller (2000) | ACCTCATAGGGGGTGCTTTT | TTGGAAGTGAAGGCTTTGAA |
| 79 | 5300 | no, smaller (4000) | CTTGAAGACCGGGTACTTCT | GCTGGCATATCTTCTCCGAC |
| 80 | 5300 | no, smaller (1000) | AACCCAGGGAGCTAAGTAGA | ATTGAATGTCCGGGGATCTT |
| 81 | 5300 | no, smaller (700) | GCCGAACAGCATATACCCT | TCAAGCGTGTTTCCTCGAT |
| 82 | 5300 | yes (5000) | TACCCGCCACTCAATTATCC | AGTATGGCGTCGAGTGTG |
| 83 | 5300 | yes (5000) | CTTTAAGCACCACGAGACGA | ACCCAAATGCAAAGCCGT |
| 84 | 5300 | yes (5000) | TAAATAAGTGCCTTCGCCCC | TGTATCTCGGCTGTCTCCA |
| 85 | 5300 | yes (5000) | CCGGAGCCTGGTAGTTCTT | TATCTTTCTCCCAGCCCGTC |
| 86 | 5300 | no, higher (7000) | CGCGCCTAGAACTATGCAA | ACACACTAGCAAGCACTGG |
| 87 | 5300 | no, higher (7000) | CCCCTCGTTTTCTACTGCTAC | GTGGGTGTGGAACTCTGTG |
| 88 | 5300 | no, higher (7000) | TAGCCTGCTTTTGTGTGGAG | CTTGCAAGTTGGTTTTGGGG |
| 89 | 5300 | yes (5000) | ACAGAAGTACAGTGAGCGT | GGGGAGGTTCAATTGGTCA |
| 90 | 5300 | no, smaller (700) | AGTCAATTCGCCTAGTACCAC | CAATTGGAGCTGCATCCTTT |
| 91 | 5300 | yes (5000) | AGGGCACTTTCCTCGAGA | ATATGCTTTGATACGGCGCT |
| 92 | 5300 | yes (5000) | GTCTTGACGCCTTGCCTAG | CACTGCATTTCAAACGCTCG |
| 93 | 5300 | no, higher (7000) | GAACACGCCGAGGATAGC | CGAAAATCTACTCAGCCTGCT |
| 94 | 5300 | yes (5000) | GCCTTCCACACGCATCTG | GATGCCCGCACTGAGAGA |
| 95 | 5300 | yes (5000) | ACCCAAGTTCTTGTCGCC | GCGGCGCATCAACTAATG |
| 96 | 5300 | no, smaller (3000) | TCCGCTGGAGAAATTGCAT | AATCTAAAAGGGGCTGCCA |
| 97 | 5300 | yes (5000) | TAAGTCACAACCCTACAGCA | TGACAGCAGTTGGGATCAA |
| 98 | 5300 | yes (5000) | CGCTTACACTGTATTTGCCC | GCCACCGTCTCTACTTGC |
| 99 | 4200 | no, smaller (600) | CTAACACGTTTCCGCCAAGT | TTCGAGGGTGTGGGTGTATT |
| 100 | 4200 | yes (4000) | TCAAGTCCCAGATGCACTCA | ATGTGGAACTTGGAGGATGC |
| 101 | 4200 | no, higher (5000) | TCAAGTCCCAGATGCACTCA | ATGTGGAACTTGGAGGATGC |
| 102 | 4200 | no, higher (5000) | TCGAAGCCATTGCTATTTTTG | TGACACCTATTCCTCAGACTCG |
| 103 | 4200 | no, smaller (400) | CAATCGCCTAGATCGCTTGT | AGTAGCAGGTCGCCTTGAAA |
| 104 | 2900 | no, smaller (1500) | CTCCCAAGGATTCTGTCCAA | AATGTTTCCCAAAGCTGACG |
| 105 | 2900 | no, higher (4000) | GGGTCTGAAAGCAGCTATGG | CATTGTTCTTGGCTGACGAA |

| | | | | |
|---|---|---|---|---|
| 106 | 2900 | no, smaller (2000) | TCAACGCTGAAAAGTATGCAA | GCAGATGATGTTGGCTTGAA |
| 107 | 2900 | no, smaller (2000) | AGCTTTAGCCACAGCCACAT | GAGAGGCACGCAGGTAAGAC |
| 108 | 2900 | no, smaller (2000) | AGCTTTAGCCACAGCCACAT | GAGAGGCACGCAGGTAAGAC |
| 109 | 2900 | yes (3000) | AGCTTTAGCCACAGCCACAT | GAGAGGCACGCAGGTAAGAC |
| 110 | 2900 | no, smaller (1500) | CAAAGGCAGGGCTAACAAAA | CACAAGTGGGAGCATCAACA |
| 111 | 1400 | yes (1750) | ACCCATCAGACTTCCACGAC | GGAATCGCCTACAGAAGCTG |
| 112 | 1400 | yes (1750) | TCGATTTGGCTGGAAACTCT | ATGCTGAACACGATGTGGAA |
| 113 | 1400 | yes (1750) | TCGATTTGGCTGGAAACTCT | ATGCTGAACACGATGTGGAA |
| 114 | 1400 | yes (1750) | GGGTACAATCAAATCGAGCTTC | GCGAAAACTGCACTCAATCA |
| 115 | 1400 | no, higher (2000) | CTTCGTTTCATTTGGCCATT | TGTGCCAAAAACACAGGCTA |
| 116 | 2100 | no, smaller (500) | GGCTACGACATTTCCGTTGT | AACCTATTCCTTGCGGACCT |
| 117 | 2100 | no, smaller (500) | GGCTACGACATTTCCGTTGT | AACCTATTCCTTGCGGACCT |
| 118 | 2100 | no, smaller (500) | TTCAATACGGATTTGCCACA | GCAAAAATAAGGGCCATCCT |
| 119 | 2100 | no, smaller (500) | TTCAATACGGATTTGCCACA | GCAAAAATAAGGGCCATCCT |
| 120 | 2100 | no, smaller (1250) | GTTGAGCAAACAGACCCACA | GGAGCCTCATAATCCGGTCT |
| 121 | 2100 | no, smaller (500) | AACTCGAATCTGGCTCGAAA | AGTGGCCTTATCGATTGGAA |
| 122 | 2100 | no, smaller (500) | GATGTTTCGTGTGGCTGTTG | GCAGTCGCTGCAGTTTGATA |
| 123 | 2100 | no, smaller (500) | GATGTTTCGTGTGGCTGTTG | GCAGTCGCTGCAGTTTGATA |
| 124 | 2100 | no, smaller (500) | TGCACATGACTGGATTCACA | CACACGAACATTGCTCCGA |
| 125 | 2100 | no, smaller (700) | TTAGAAAGCAAGTACCGGCA | TTCTGCCATCGTGTTGCC |
| 126 | 2100 | yes (2000) | GTGGGGCCTCATAGATACGT | TATGCGCTGAGGTACACTTG |
| 127 | 2100 | no, smaller (700) | CCGGCCCATGTTAAGCTTTA | GAGGCAGCGGATCAATTCA |
| 128 | 2100 | no, smaller (500) | TGTTCGGGTGGTAAAAGCG | CAACGTTCCCAGGACACC |
| 129 | 2100 | no, smaller (700) | ACGCCTCGGATTTGACATC | CCGTTGGCATTTTGTGGATA |
| 130 | 2100 | no, smaller (500) | ATAGCACATTCAGCCACACG | TGCTGAATTCGGAAAGAGCT |
| 131 | 2100 | yes (2000) | GGTTTCGATTCGGTATTGGTTG | ATTGCTTCGTGTTAGGACCC |
| 132 | 2100 | no, smaller (500) | CTGAGTCGAGCTGGTAGGT | CGACATTTTCTGCGGCCG |
