## Supplementary material for "Effects of *Wolbachia* on transposable element activity largely depend on *Drosophila melanogaster* host genotype": Table S2 - available upon acceptance

| Supplementary Table S2 | | | | | |
|---|---|---|---|---|---|
| **TE expression data** | | | | | |
| **DGRP** | **TE** | **Wolbachia status** | **Replicate** | **TE Cq** | **EF1 Cq** |

Data available upon acceptance for publication
