## Supplementary material for "Effects of *Wolbachia* on transposable element activity largely depend on *Drosophila melanogaster* host genotype": Table S3

| Supplementary Table S3 | | | | |
|---|---|---|---|---|
| **Statistical significance of TE expression differences with Wolbachia** | | | | |
| **TE** | **DGRP** | **Degrees of freedom (df)** | **T-test (t)** | **Statistical differences (p-value)** |
| *412* | RAL-136 | 6.8911 | 1,192 | 0,2725 |
| *412* | RAL-142 | 7.0000 | 2,431 | 0,0453 * |
| *412* | RAL-149 | 5.4629 | 3,378 | 0,0172 * |
| *412* | RAL-181 | 7.0000 | 3,255 | 0,01396 * |
| *412* | RAL-21 | 6.2841 | -3,142 | 0,01883 * |
| *412* | RAL-237 | 14.0000 | 0,965 | 0,351 |
| *412* | RAL-280 | 7.0000 | 1,269 | 0,245 |
| *412* | RAL-320 | 14.0000 | 1,611 | 0,1295 |
| *412* | RAL-321 | 6.99988 | -0,4 | 0,701 |
| *412* | RAL-338 | 7.00000 | -0,108 | 0,917 |
| *412* | RAL-352 | 14.0000 | 1,869 | 0,0827 . |
| *412* | RAL-370 | 7.0000 | -1,05 | 0,329 |
| *412* | RAL-374 | 14.00000 | 0,463 | 0,651 |
| *412* | RAL-440 | 6.8022 | -2,146 | 0,0701 . |
| *412* | RAL-441 | 14.0000 | 1,096 | 0,291 |
| *412* | RAL-535 | 7.0000 | 1,91 | 0,0977 . |
| *412* | RAL-595 | 14.0000 | 0,867 | 0,401 |
| *412* | RAL-69 | 7.0000 | 0,737 | 0,485 |
| *412* | RAL-712 | 14.0000 | -2,743 | 0,01587 * |
| *412* | RAL-737 | 14.0000 | 3,624 | 0,002762 ** |
| *412* | RAL-738 | 7.0000 | 2,031 | 0,0818 . |
| *412* | RAL-801 | 7.00000 | 0,34 | 0,744 |
| *412* | RAL-820 | 7.0000 | -1,985 | 0,0875 . |
| *412* | RAL-855 | 7.0000 | 1,027 | 0,339 |
| *412* | RAL-882 | 7.0000 | 0,922 | 0,387 |
| *1360* | RAL-136 | 12.0000 | 0,74 | 0,474 |
| *1360* | RAL-142 | 7.00000 | 0,257 | 0,804 |
| *1360* | RAL-149 | 11.00000 | 1,053 | 0,315 |
| *1360* | RAL-181 | 7.00000 | 5,586 | 0,000828 *** |
| *1360* | RAL-21 | 7.0000 | -4,067 | 0,00477 ** |
| *1360* | RAL-237 | 7.0000 | -0,861 | 0,418 |
| *1360* | RAL-280 | 12.0000 | 0,793 | 0,443 |
| *1360* | RAL-320 | 14.0000 | 0,752 | 0,464 |
| *1360* | RAL-321 | 12.0000 | -1,192 | 0,256 |
| *1360* | RAL-338 | 6.4694 | 2,344 | 0,0544 . |
| *1360* | RAL-352 | 7.0000 | 1,996 | 0,0827 . |
| *1360* | RAL-370 | 5.6893 | -1,145 | 0,298 |
| *1360* | RAL-374 | 14.00000 | -1,502 | 0,1553 |
| *1360* | RAL-440 | 14.0000 | -0,615 | 0,548 |
| *1360* | RAL-441 | 14.00000 | 0,212 | 0,835 |
| *1360* | RAL-535 | 7.000000 | 0,097 | 0,925 |
| *1360* | RAL-595 | 14.0000 | -0,353 | 0,73 |
| *1360* | RAL-69 | 6.90130 | -0,108 | 0,917 |
| *1360* | RAL-712 | 14.0000 | -3,019 | 0,009203 ** |
| *1360* | RAL-737 | 7.00000 | -6,067 | 0,000507 *** |
| *1360* | RAL-738 | 7.0000 | 1,664 | 0,1401 |
| *1360* | RAL-801 | 7.0000 | -1,529 | 0,17 |
| *1360* | RAL-820 | 14.0000 | -1,954 | 0,0709 . |
| *1360* | RAL-855 | 7.00000 | 0,459 | 0,66 |
| *1360* | RAL-882 | 7.0000 | 1,532 | 0,1693 |
| *blastopia* | RAL-136 | 6.6966 | -0,898 | 0,4 |
| *blastopia* | RAL-142 | 6.16874 | 0,253 | 0,808 |

| | | | | |
|---|---|---|---|---|
| *blastopia* | RAL-149 | 6.000000 | -1,771 | 0,127 |
| *blastopia* | RAL-181 | 7.00000 | 5,985 | 0,00055 *** |
| *blastopia* | RAL-21 | 7.0000 | -2,196 | 0,0641 . |
| *blastopia* | RAL-237 | 7.0000 | -0,988 | 0,356 |
| *blastopia* | RAL-280 | 14.00000 | 0,261 | 0,798 |
| *blastopia* | RAL-320 | 14.00000 | -0,161 | 0,874 |
| *blastopia* | RAL-321 | 14.0000 | -3,437 | 0,004009 ** |
| *blastopia* | RAL-338 | 7.14843 | -3,537 | 0,000194 *** |
| *blastopia* | RAL-352 | 13.00000 | 0,325 | 0,75 |
| *blastopia* | RAL-370 | 7.0000 | -0,606 | 0,564 |
| *blastopia* | RAL-374 | 14.000000 | 0,006 | 0,995 |
| *blastopia* | RAL-440 | 6.6346 | -2,666 | 0,0338* |
| *blastopia* | RAL-441 | 7.0000 | 1,138 | 0,293 |
| *blastopia* | RAL-535 | 7.00000 | 0,552 | 0,598 |
| *blastopia* | RAL-595 | 13.0000 | -1,894 | 0,0807 . |
| *blastopia* | RAL-69 | 7.00000 | -6,28 | 0,000412 *** |
| *blastopia* | RAL-712 | 7.0000 | -2,806 | 0,02628 * |
| *blastopia* | RAL-737 | 7.00000 | -0,495 | 0,635 |
| *blastopia* | RAL-738 | 14.00000 | -0,111 | 0,913 |
| *blastopia* | RAL-801 | 13.0000 | -0,873 | 0,399 |
| *blastopia* | RAL-820 | 14.00000 | 0,413 | 0,686 |
| *blastopia* | RAL-855 | 7.0000 | 1,007 | 0,348 |
| *blastopia* | RAL-882 | 7.0000 | -1,17 | 0,28 |
| *blood* | RAL-136 | 4.5202 | 1,367 | 0,2357 |
| *blood* | RAL-142 | 7.0000 | 2,541 | 0,03858 * |
| *blood* | RAL-149 | 6.0000 | 3,065 | 0,0221 * |
| *blood* | RAL-181 | 7.0000 | 2,4 | 0,0475 * |
| *blood* | RAL-21 | 7.0000 | -4,697 | 0,002217 ** |
| *blood* | RAL-237 | 14.0000 | 1,953 | 0,0712 . |
| *blood* | RAL-280 | 7.0000 | 0,95 | 0,374 |
| *blood* | RAL-320 | 14.0000 | 1,891 | 0,0795 . |
| *blood* | RAL-321 | 14.0000 | -2,504 | 0,02528 * |
| *blood* | RAL-338 | 13.0000 | -0,859 | 0,406 |
| *blood* | RAL-352 | 14.0000 | 0,756 | 0,462 |
| *blood* | RAL-370 | 7.0000 | -2,09 | 0,075 . |
| *blood* | RAL-374 | 14.0000 | -1,392 | 0,1858 |
| *blood* | RAL-440 | 5.0453 | -3,126 | 0,0258 * |
| *blood* | RAL-441 | 14.0000 | 0,618 | 0,546 |
| *blood* | RAL-535 | 6.99998 | 0,377 | 0,717 |
| *blood* | RAL-595 | 14.0000 | 5,549 | 7 .17e-05 *** |
| *blood* | RAL-69 | 6.0259 | 0,989 | 0,361 |
| *blood* | RAL-712 | 13.0000 | -3,056 | 0,009185 ** |
| *blood* | RAL-737 | 6.0000 | 0,822 | 0,443 |
| *blood* | RAL-738 | 7.0000 | 1,663 | 0,1404 |
| *blood* | RAL-801 | 7.0000 | -0,384 | 0,712 |
| *blood* | RAL-820 | 14.0000 | -0,81 | 0,431 |
| *blood* | RAL-855 | 14.0000 | 1,885 | 0,0804 . |
| *blood* | RAL-882 | 14.0000 | 2,909 | 0,01144 * |
| *copia* | RAL-136 | 6.998364 | -0,42 | 0,686 |
| *copia* | RAL-142 | 7.0000 | 2,512 | 0,0403 * |
| *copia* | RAL-149 | 6.0000 | 2,724 | 0,0345 * |
| *copia* | RAL-181 | 7.0000 | 2,121 | 0,0716 . |
| *copia* | RAL-21 | 13.0000 | 0,647 | 0,529 |
| *copia* | RAL-237 | 7.00000 | 0,016 | 0,988 |
| *copia* | RAL-280 | 6.1421 | -0,043 | 0,967 |

| | | | | |
|---|---|---|---|---|
| *copia* | RAL-320 | 14.0000 | 0,943 | 0,362 |
| *copia* | RAL-321 | 14.0000 | 1,032 | 0,32 |
| *copia* | RAL-338 | 7.0000 | -3,47 | 0,0104 * |
| *copia* | RAL-352 | 14.0000 | 0,958 | 0,354 |
| *copia* | RAL-370 | 7.0000 | -1,058 | 0,325 |
| *copia* | RAL-374 | 13.0000 | -1,336 | 0,2043 |
| *copia* | RAL-440 | 6.1544 | -5,771 | 0,00108 ** |
| *copia* | RAL-441 | 14.0000 | -0,846 | 0,412 |
| *copia* | RAL-535 | 14.00000 | -0,524 | 0,608 |
| *copia* | RAL-595 | 14.0000 | 1,211 | 0,246 |
| *copia* | RAL-69 | 7.0000 | 0,848 | 0,425 |
| *copia* | RAL-712 | 14.0000 | -3,746 | 0,002170 ** |
| *copia* | RAL-737 | 7.0000 | 4,469 | 0,002905 ** |
| *copia* | RAL-738 | 7.0000 | 1,329 | 0,226 |
| *copia* | RAL-801 | 7.00000 | 0,28 | 0,787 |
| *copia* | RAL-820 | 7.0021 | -2,241 | 0,0600 . |
| *copia* | RAL-855 | 14.0000 | 2,289 | 0,03810 * |
| *copia* | RAL-882 | 7.0000 | 0,67 | 0,524 |
| *Cr1a* | RAL-136 | 13.0000 | -2,454 | 0,02898 * |
| *Cr1a* | RAL-142 | 7.00000 | 2,271 | 0,05738 . |
| *Cr1a* | RAL-149 | 6.00000 | 1,297 | 0,242 |
| *Cr1a* | RAL-181 | 14.00000 | 5,64 | 6,10e-05 *** |
| *Cr1a* | RAL-21 | 14.0000 | -2,524 | 0,02432 * |
| *Cr1a* | RAL-237 | 7.0000 | -1,547 | 0,166 |
| *Cr1a* | RAL-280 | 7.0000 | 0,682 | 0,517 |
| *Cr1a* | RAL-320 | 14.0000 | -0,514 | 0,616 |
| *Cr1a* | RAL-321 | 14.0000 | -3,43 | 0,004059 ** |
| *Cr1a* | RAL-338 | 7.00000 | 2,913 | 0,02256 * |
| *Cr1a* | RAL-352 | 7.00000 | 1,407 | 0,202 |
| *Cr1a* | RAL-370 | 13.0000 | -0,961 | 0,354 |
| *Cr1a* | RAL-374 | 6.6513 | 0,843 | 0,428 |
| *Cr1a* | RAL-440 | 6.8131 | -1,949 | 0,0935 . |
| *Cr1a* | RAL-441 | 14.00000 | 0,207 | 0,839 |
| *Cr1a* | RAL-535 | 7.00000 | 1,069 | 0,321 |
| *Cr1a* | RAL-595 | 14.0000 | -1,803 | 0,0929 . |
| *Cr1a* | RAL-69 | 7.00000 | 0,227 | 0,827 |
| *Cr1a* | RAL-712 | 14.0000 | -2,612 | 0,02051 * |
| *Cr1a* | RAL-737 | 7.0000 | -0,558 | 0,594 |
| *Cr1a* | RAL-738 | 13.0000 | -0,514 | 0,616 |
| *Cr1a* | RAL-801 | 14.0000 | -0,526 | 0,607 |
| *Cr1a* | RAL-820 | 14.0000 | -2,108 | 0,05352 . |
| *Cr1a* | RAL-855 | 7.0000 | -0,522 | 0,618 |
| *Cr1a* | RAL-882 | 7.00000 | -0,366 | 0,725 |
| *gypsy5* | RAL-136 | 4.02955 | -0,102 | 0,923 |
| *gypsy5* | RAL-142 | 7.00000 | 2,381 | 0,0488 * |
| *gypsy5* | RAL-149 | 6.00000 | 0,437 | 0,6775 |
| *gypsy5* | RAL-181 | 7.00000 | 5,483 | 0,000923 *** |
| *gypsy5* | RAL-21 | 6.6635 | -3,722 | 0,00813 ** |
| *gypsy5* | RAL-237 | 7.000000 | 0,004 | 0,997 |
| *gypsy5* | RAL-280 | 6.9998 | 3,52 | 0,009729 ** |
| *gypsy5* | RAL-320 | 13.0000 | -1,702 | 0,1126 |
| *gypsy5* | RAL-321 | 13.0000 | -1,942 | 0,0741 . |
| *gypsy5* | RAL-338 | 7.0000 | -5,805 | 0,00066 *** |
| *gypsy5* | RAL-352 | 7.00000 | 1,846 | 0,1074 |
| *gypsy5* | RAL-370 | 13.00000 | 0,622 | 0,545 |

| | | | | |
|---|---|---|---|---|
| *gypsy5* | RAL-374 | 14.00000 | -0,516 | 0,614 |
| *gypsy5* | RAL-440 | 6.3836 | -1,768 | 0,125 |
| *gypsy5* | RAL-441 | 14.00000 | 0,376 | 0,712 |
| *gypsy5* | RAL-535 | 6.9546 | -0,762 | 0,471 |
| *gypsy5* | RAL-595 | 14.0000 | -0,959 | 0,354 |
| *gypsy5* | RAL-69 | 7.0000 | 1,096 | 0,309 |
| *gypsy5* | RAL-712 | 7.0000 | -1,221 | 0,262 |
| *gypsy5* | RAL-737 | 7.0000 | -4,117 | 0,00448 ** |
| *gypsy5* | RAL-738 | 13.0000 | -0,458 | 0,654 |
| *gypsy5* | RAL-801 | 13.0000000 | -0,002 | 0,999 |
| *gypsy5* | RAL-820 | 13.0000 | 0,928 | 0,37 |
| *gypsy5* | RAL-855 | 14.00000 | 1,006 | 0,331 |
| *gypsy5* | RAL-882 | 7.0000 | 0,896 | 0,4 |
| *Idefix* | RAL-136 | 7.00000 | 0,697 | 0,508 |
| *Idefix* | RAL-142 | 7.0000 | 1,234 | 0,257 |
| *Idefix* | RAL-149 | 7.0000 | 1,486 | 0,181 |
| *Idefix* | RAL-181 | 7.0281 | 3,213 | 0,014723 * |
| *Idefix* | RAL-21 | 6.9155 | -3,263 | 0,01404 * |
| *Idefix* | RAL-237 | 7.00000 | -0,205 | 0,844 |
| *Idefix* | RAL-280 | 14.00000 | 0,219 | 0,829 |
| *Idefix* | RAL-320 | 14.00000 | 0,339 | 0,74 |
| *Idefix* | RAL-321 | 5.85120 | -3,913 | 0,00827 ** |
| *Idefix* | RAL-338 | 7.0000 | -2,206 | 0,0632 . |
| *Idefix* | RAL-352 | 14.00000 | 0,297 | 0,771 |
| *Idefix* | RAL-370 | 7.0000 | 1,984 | 0,0876 . |
| *Idefix* | RAL-374 | 14.00000 | -0,095 | 0,926 |
| *Idefix* | RAL-440 | 7.0102 | -1,632 | 0,1467 |
| *Idefix* | RAL-441 | 7.0000 | 1,77 | 0,1199 |
| *Idefix* | RAL-535 | 14.00000 | 0,315 | 0,757 |
| *Idefix* | RAL-595 | 10.00000 | -0,057 | 0,956 |
| *Idefix* | RAL-69 | 7.0000 | 1,896 | 0,0998 . |
| *Idefix* | RAL-712 | 7.0000 | -1,34 | 0,222 |
| *Idefix* | RAL-737 | 7.0000 | -4,77 | 0,002035 ** |
| *Idefix* | RAL-738 | 14.00000 | -0,143 | 0,889 |
| *Idefix* | RAL-801 | 7.00000 | 0,153 | 0,883 |
| *Idefix* | RAL-820 | 6.4753 | 3,383 | 0,0132 * |
| *Idefix* | RAL-855 | 6.9979 | 1,539 | 0,1676 |
| *Idefix* | RAL-882 | 6.7860 | 1,535 | 0,17 |
| *Juan* | RAL-136 | 7.0000 | -0,091 | 0,93 |
| *Juan* | RAL-142 | 5.75776 | 0,348 | 0,74 |
| *Juan* | RAL-149 | 6.9589 | 3,233 | 0,01452 * |
| *Juan* | RAL-181 | 7.0000 | 0,648 | 0,538 |
| *Juan* | RAL-21 | 7.0000 | -4,021 | 0,00505 ** |
| *Juan* | RAL-237 | 7.00000 | 0,355 | 0,733 |
| *Juan* | RAL-280 | 14.00000 | -0,046 | 0,964 |
| *Juan* | RAL-320 | 14.0000 | 0,373 | 0,715 |
| *Juan* | RAL-321 | 4.90491 | 0,999 | 0,364 |
| *Juan* | RAL-338 | 7.0000 | -0,453 | 0,664 |
| *Juan* | RAL-352 | 14.00000 | 0,421 | 0,68 |
| *Juan* | RAL-370 | 7.0000 | -1,671 | 0,139 |
| *Juan* | RAL-374 | 14.00000 | -1,785 | 0,0959 . |
| *Juan* | RAL-440 | 6.7921 | -3,151 | 0,01677 * |
| *Juan* | RAL-441 | 14.0000 | 0,554 | 0,588 |
| *Juan* | RAL-535 | 14.00000 | -0,32 | 0,754 |
| *Juan* | RAL-595 | 14.00000 | 0,434 | 0,671 |

| | | | | |
|---|---|---|---|---|
| *Juan* | RAL-69 | 7.0000 | 0,578 | 0,581 |
| *Juan* | RAL-712 | 7.0000 | -5,073 | 0,00144 ** |
| *Juan* | RAL-737 | 7.0000 | -0,753 | 0,476 |
| *Juan* | RAL-738 | 7.0000 | 0,489 | 0,64 |
| *Juan* | RAL-801 | 7.0000 | -1,636 | 0,146 |
| *Juan* | RAL-820 | 6.63341 | -0,32 | 0,759 |
| *Juan* | RAL-855 | 7.0000 | 0,903 | 0,397 |
| *Juan* | RAL-882 | 7.00000 | 0,258 | 0,804 |
| *mdg1* | RAL-136 | 7.000000 | -0,03 | 0,977 |
| *mdg1* | RAL-142 | 6.6493 | 1,707 | 0,1339 |
| *mdg1* | RAL-149 | 7.0000 | 1,295 | 0,2363 |
| *mdg1* | RAL-181 | 7.00000 | 4,065 | 0,00478 ** |
| *mdg1* | RAL-21 | 7.0000 | -4,758 | 0,002065 ** |
| *mdg1* | RAL-237 | 14.0000 | -0,651 | 0,526 |
| *mdg1* | RAL-280 | 7.00000 | 0,292 | 0,779 |
| *mdg1* | RAL-320 | 14.0000 | 0,323 | 0,752 |
| *mdg1* | RAL-321 | 14.0000 | -3,263 | 0,005666 ** |
| *mdg1* | RAL-338 | 7.0000 | -0,084 | 0,936 |
| *mdg1* | RAL-352 | 14.0000 | -0,647 | 0,528 |
| *mdg1* | RAL-370 | 7.0000 | -3,147 | 0,0162 * |
| *mdg1* | RAL-374 | 14.00000 | 1,755 | 0,1011 |
| *mdg1* | RAL-440 | 6.5603 | -3,76 | 0,00797 ** |
| *mdg1* | RAL-441 | 14.0000 | 1,265 | 0,2267 |
| *mdg1* | RAL-535 | 6.99999 | 0,152 | 0,884 |
| *mdg1* | RAL-595 | 14.0000 | 0,77 | 0,454 |
| *mdg1* | RAL-69 | 5.4550 | 1,192 | 0,283 |
| *mdg1* | RAL-712 | 13.0000 | -2,767 | 0,01600 * |
| *mdg1* | RAL-737 | 6.8714 | -0,863 | 0,417 |
| *mdg1* | RAL-738 | 14.0000 | 2,463 | 0,02736 * |
| *mdg1* | RAL-801 | 7.0000 | -2,311 | 0,0541 . |
| *mdg1* | RAL-820 | 12.0000 | -3,513 | 0,004274 ** |
| *mdg1* | RAL-855 | 7.0000 | 1,469 | 0,1852 |
| *mdg1* | RAL-882 | 7.0000 | 0,819 | 0,44 |
| *opus* | RAL-136 | 7.00000 | -0,384 | 0,712 |
| *opus* | RAL-142 | 13.00000 | 2,39 | 0,03269 * |
| *opus* | RAL-149 | 7.00000 | -0,766 | 0,468 |
| *opus* | RAL-181 | 7.00000 | -1,751 | 0,1233 |
| *opus* | RAL-21 | 6.8085 | -2,994 | 0,02078 * |
| *opus* | RAL-237 | 7.0000 | -0,902 | 0,397 |
| *opus* | RAL-280 | 14.0000 | 3,379 | 0,004501 ** |
| *opus* | RAL-320 | 7.0000 | 2,781 | 0,02726 * |
| *opus* | RAL-321 | 14.0000 | -3,985 | 0,00136 ** |
| *opus* | RAL-338 | 7.0000 | -1,023 | 0,34 |
| *opus* | RAL-352 | 7.0000 | -0,848 | 0,424 |
| *opus* | RAL-370 | 7.0000 | -1,058 | 0,325 |
| *opus* | RAL-374 | 6.9131 | -1,155 | 0,286 |
| *opus* | RAL-440 | 7.0000 | -1,223 | 0,261 |
| *opus* | RAL-441 | 7.00000 | 1,298 | 0,235 |
| *opus* | RAL-535 | 7.00000 | -0,46 | 0,659 |
| *opus* | RAL-595 | 14.0000 | -2,178 | 0,04701 * |
| *opus* | RAL-69 | 7.00000 | -0,439 | 0,674 |
| *opus* | RAL-712 | 7.0000 | -1,951 | 0,0920 . |
| *opus* | RAL-737 | 7.00000 | 0,644 | 0,54 |
| *opus* | RAL-738 | 14.0000 | -0,564 | 0,582 |
| *opus* | RAL-801 | 14.0000 | -0,431 | 0,673 |

| | | | | |
|---|---|---|---|---|
| *opus* | RAL-820 | 13.0000 | -1,21 | 0,248 |
| *opus* | RAL-855 | 7.00000 | -0,432 | 0,679 |
| *opus* | RAL-882 | 7.0000 | -1,116 | 0,301 |
| *P-element* | RAL-136 | 7.0000 | 1,809 | 0,1134 |
| *P-element* | RAL-142 | 7.00000 | 0,314 | 0,762 |
| *P-element* | RAL-149 | 6.9999 | 2,039 | 0,0808 . |
| *P-element* | RAL-181 | 7.0000 | 3,281 | 0,01347 * |
| *P-element* | RAL-21 | 14.0000 | -4,721 | 0,000328 *** |
| *P-element* | RAL-237 | 14.0000 | 0,554 | 0,588 |
| *P-element* | RAL-280 | 7.0000 | 1,521 | 0,172 |
| *P-element* | RAL-320 | 14.00000 | -0,172 | 0,866 |
| *P-element* | RAL-321 | 14.00000 | -3,748 | 0,002163 ** |
| *P-element* | RAL-338 | 7.0000 | 1,683 | 0,1363 |
| *P-element* | RAL-352 | 7.0000 | 1,306 | 0,233 |
| *P-element* | RAL-370 | 7.0000 | -2,423 | 0,0459 * |
| *P-element* | RAL-374 | 14.0000 | -0,957 | 0,355 |
| *P-element* | RAL-440 | 5.8482 | -2,082 | 0,08366 . |
| *P-element* | RAL-441 | 14.0000 | 0,781 | 0,448 |
| *P-element* | RAL-535 | 14.0000 | 4,583 | 0,000426 *** |
| *P-element* | RAL-595 | 14.0000 | -0,386 | 0,705 |
| *P-element* | RAL-69 | 6.99972 | 0,302 | 0,772 |
| *P-element* | RAL-712 | 14.0000 | -5,025 | 0,000186 *** |
| *P-element* | RAL-737 | 7.0000 | -1,435 | 0,195 |
| *P-element* | RAL-738 | 7.0000 | 1,334 | 0,224 |
| *P-element* | RAL-801 | 7.0000 | -1,388 | 0,208 |
| *P-element* | RAL-820 | 6.8606 | -2,258 | 0,0592 . |
| *P-element* | RAL-855 | 14.0000 | 2,865 | 0,01247 * |
| *P-element* | RAL-882 | 7.0000 | 0,994 | 0,353 |
| *pogo* | RAL-136 | 7.0000 | 1,171 | 0,28 |
| *pogo* | RAL-142 | 6.91144 | 0,208 | 0,841 |
| *pogo* | RAL-149 | 7.0000 | 1,435 | 0,194 |
| *pogo* | RAL-181 | 7.00000 | 2,294 | 0,0555 . |
| *pogo* | RAL-21 | 7.0000 | -5,395 | 0,00101 ** |
| *pogo* | RAL-237 | 14.00000 | 0,04 | 0,969 |
| *pogo* | RAL-280 | 7.0000 | 1,091 | 0,311 |
| *pogo* | RAL-320 | 14.0000 | 0,463 | 0,65 |
| *pogo* | RAL-321 | 6.9627 | -1,546 | 0,1662 |
| *pogo* | RAL-338 | 7.0000 | -1,548 | 0,166 |
| *pogo* | RAL-352 | 14.0000 | 0,954 | 0,356 |
| *pogo* | RAL-370 | 7.00000 | -3,622 | 0,00848 ** |
| *pogo* | RAL-374 | 14.0000 | -2,559 | 0,02273 * |
| *pogo* | RAL-440 | 6.0000 | -1,356 | 0,224 |
| *pogo* | RAL-441 | 14.0000 | 0,451 | 0,659 |
| *pogo* | RAL-535 | 7.0000 | -0,782 | 0,46 |
| *pogo* | RAL-595 | 7.0000 | -1,038 | 0,334 |
| *pogo* | RAL-69 | 7.00000 | 0,183 | 0,86 |
| *pogo* | RAL-712 | 14.0000 | -2,919 | 0,01122 * |
| *pogo* | RAL-737 | 7.0000 | 0,898 | 0,399 |
| *pogo* | RAL-738 | 14.0000 | 1,587 | 0,1349 |
| *pogo* | RAL-801 | 7.0000 | -1,008 | 0,347 |
| *pogo* | RAL-820 | 13.0000 | -1,821 | 0,0916 . |
| *pogo* | RAL-855 | 14.00000 | 1,893 | 0,0792 . |
| *pogo* | RAL-882 | 7.00000 | 0,237 | 0,819 |
| *Quasimodo* | RAL-136 | 7.0000 | -4,257 | 0,003762 ** |
| *Quasimodo* | RAL-142 | 4.4967 | 3,011 | 0,0340 * |

| | | | | |
|---|---|---|---|---|
| *Quasimodo* | RAL-149 | 7.0000 | 0,943 | 0,377 |
| *Quasimodo* | RAL-181 | 7.00000 | 0,546 | 0,602 |
| *Quasimodo* | RAL-21 | 7.0000 | -4,063 | 0,004794 ** |
| *Quasimodo* | RAL-237 | 7.0000 | -2,338 | 0,0520 . |
| *Quasimodo* | RAL-280 | 14.00000 | 0,451 | 0,659 |
| *Quasimodo* | RAL-320 | 14.0000 | 1,658 | 0,1195 |
| *Quasimodo* | RAL-321 | 14.0000 | -3,325 | 0,00501 ** |
| *Quasimodo* | RAL-338 | 7.00000 | 4,218 | 0,00395 ** |
| *Quasimodo* | RAL-352 | 14.00000 | 0,327 | 0,749 |
| *Quasimodo* | RAL-370 | 7.00000 | -0,608 | 0,562 |
| *Quasimodo* | RAL-374 | 7.0000 | -0,927 | 0,385 |
| *Quasimodo* | RAL-440 | 7.0000 | -2,193 | 0,0644 . |
| *Quasimodo* | RAL-441 | 7.0000 | 1,478 | 0,1829 |
| *Quasimodo* | RAL-535 | 14.00000 | -0,455 | 0,656 |
| *Quasimodo* | RAL-595 | 14.0000 | -2,833 | 0,0133 * |
| *Quasimodo* | RAL-69 | 6.96916 | 0,598 | 0,569 |
| *Quasimodo* | RAL-712 | 14.0000 | -1,004 | 0,333 |
| *Quasimodo* | RAL-737 | 7.0000 | -1,647 | 0,144 |
| *Quasimodo* | RAL-738 | 14.000000 | 0,034 | 0,973 |
| *Quasimodo* | RAL-801 | 14.00000 | 0,198 | 0,846 |
| *Quasimodo* | RAL-820 | 13.00000 | 0,017 | 0,986 |
| *Quasimodo* | RAL-855 | 14.00000 | 0,304 | 0,765 |
| *Quasimodo* | RAL-882 | 7.00000 | -0,08 | 0,939 |
