## Supplementary material for "Effects of *Wolbachia* on transposable element activity largely depend on *Drosophila melanogaster* host genotype": Table S4

| Supplementary Table S4 | | | | |
|---|---|---|---|---|
| **Primers** | | | | |
| **TE** | **Forward primer (5' – 3')** | **Reverse primer (5' – 3')** | **Reference** | **Efficiency (%)** |
| *412* | TCATAACGGACATGGGAACA | CCAACTGTCTGGTGGTGATG | designed in Primer3 | 100,67748 |
| *1360* | TCTAGCACAACACGCACACT | GTGACGGCCAAAATTGCTGT | designed in Primer3 | 93,28801353 |
| *blastopia* | GAGCAGTCAATCGTCCGTAA | TCTATAGTCCACGCAAACGC | Chen et al. 2016 | 98,65437494 |
| *blood* | AACAATAGAAAGAAGCCACCGAA | AGTCATGGACTATTGAGGGTGTT | Handler et al. 2011 | 96,64238427 |
| *copia* | TGCCAGAGAGCAAGTTCAGA | GCAAACCCAATTTGTCTCGT | designed in Primer3 | 100,2605663 |
| *Cr1a* | TGGCCGTACAAGTGATGACC | TCATCTCGTTCGCAACCACA | designed in Primer3 | 97,33446929 |
| *gypsy 5* | GCCCAGAGACAACGACAGAA | CTGTCTTTGCTGTCCCGGAT | designed in Primer3 | 95,32455343 |
| *H-element* | CATTAAGTCGGAAGGCCAAA | CTTGCTCTTCCGCTATCCAC | designed in Primer3 | (not used in qPCR) |
| *Idefix* | CGCTCTAGTGGACAGAACCA | TGAAATGAGGCATTTGGGTA | Chen et al. 2016 | 104,9522518 |
| *jockey* | GCGGATTAACAAGGGGCTCT | CCTGGGAGATAGATGCGCTG | designed in Primer3 | (not used in qPCR) |
| *Juan* | GGGGCAAAATTCTCAATGAA | GCGGAATATATGTGGGTTGC | designed in Primer3 | 104,4958212 |
| *mdg1* | GTCAGAAGGAGGCCATTCAGGAA | GTTGCTGGCGGTTTCTGTTATTG | Navarro et al. 2009 | 91,87285071 |
| *opus* | CGAGGAGTGGGGAGAGATT | TGCGAAAATCTGCCTGAACC | Specchia et al. 2010 | 101,4399028 |
| *P-element* | TGAGTGCTCGCAACCTTATG | TTTGAAATGGGAGCCTTTTG | designed in Primer3 | 94,08570477 |
| *pogo* | CCAGCGATAACGAAGAAAGC | GCTGCAAACCCATCCTTAAA | designed in Primer3 | 105,4403078 |
| *Quasimodo* | TCTACAGTGCCATCGAGAGG | TAGTTCAGCCCAAGTGTTGC | Chen et al. 2016 | 100,8172005 |
| **Gene** | | | | |
| *actin* | GCGTCGGTCAATTCAATCTT | AAGCTGCAACCTCTTCGCA | Ponton et al. 2011 | (not used in qPCR) |
| *EF1* | GCGTGGGTTTGTGATCAGTT | GATCTTCTCCTTGCCCATCC | Ponton et al. 2011 | 98,0965242 |
| *RPL32* | ATGCTAAGCTGTCGCACAAATG | GTTCGATCCGTAACCGATGT | designed in Primer3 | (not used in qPCR) |
| *wsp* | TGGTCCAATAAGTGATGAAGAAAC | AAAAATTAAACGCTACTCCA | Teixeira et al. 2008 | (not used in qPCR) |

### Primer sequence references:


Chen H, Zheng X, Xiao D, Zheng Y. 2016. Age-associated de-repression of retrotransposons in the Drosophila fat body, its potential cause and consequence. *Aging Cell* **15**: 542-552.

Handler D, Olivieri D, Novatchkova M, Gruber FS, Meixner K, Mechtler K, Stark A, Sachidanandam R, Brennecke J. 2011. A systematic analysis of *Drosophila* TUDOR domain-containing proteins identifies Vreteno and the Tdrd12 family as essential primary piRNA pathway factors. *EMBO J* **30**: 3977-3993.

Navarro C, Bullock S, Lehmann R. 2009. Altered dynein-dependent transport in piRNA pathway mutants. *PNAS* **106**: 9691-9696.

Ponton F, Chapuis M-P, Pernice M, Sword GA, Simpson SJ. 2011.Evaluation of potential reference genes for reverse transcription-qPCR studies of physiological responses in Drosophila melanogaster. *J Insect Physiol* **57**: 840-850.

Specchia V, Piacentini L, Tritto P, Fanti L, D'Alessandro R, Palumbo G, Pimpinelli S, Bozzetti MP. 2010. Hsp90 prevents phenotypic variation by suppressing the mutagenic activity of transposons. *Nature* **463**: 662-665.

Teixeira L, Ferreira Á, Ashburner M. 2008. The bacterial symbiont *Wolbachia* induces resistance to RNA viral infections in Drosophila melanogaster. PLOS Biol 6: e1000002.
